# Chromosome-scale and haplotype-resolved genome assembly of a tetraploid potato cultivar

**DOI:** 10.1101/2021.05.15.444292

**Authors:** Hequan Sun, Wen-Biao Jiao, José A. Campoy, Kristin Krause, Manish Goel, Kat Folz-Donahue, Christian Kukat, Bruno Huettel, Korbinian Schneeberger

## Abstract

Potato is the most important tuber crop in the world. However, separate reconstruction of the four haplotypes of its autotetraploid genome remained an unsolved challenge. Here, we report the 3.1 Gb haplotype-resolved (at 99.6% precision), chromosome-scale assembly of the potato cultivar ‘Otava’ using high-quality long reads coupled with single-cell sequencing of 717 pollen genomes and Hi-C data. Unexpectedly, almost 50% of the genome were found to be identical-by-descent due to recent inbreeding, which contrasted by highly abundant structural rearrangements involving around 20% of the genome. Among 38,214 genes, only 54% were present in four haplotypes with an average of 3.2 copies per gene. Analyzing the leaf transcriptome as example, we found that 11% of the genes featured differently expressed alleles in at least one of the haplotypes, of which 25% are likely regulated through allele-specific DNA methylation. Our work sheds light on the recent breeding history of potato, the functional organization of its tetraploid genome and has the potential to strengthen the future of genomics-assisted breeding.

---

Potato *(Solanum tuberosum)* is by far the most important tuber crop and is among the five most produced crops in the world. Globally more than 350 billion kilograms of potato are produced per year with an increasing trend particularly in developing countries in Asia^1^. Despite the social and economic importance, the breeding success of potato remained low over the past decades due to its heterozygous, autotetraploid genome and the high levels of inbreeding depression, which challenge usual breeding strategies commonly applied to inbred, diploid crops^2,3^.

A fundamental tool for modern breeding is the availability of reference sequences. The reference sequence for potato was generated from a double haploid plant, *DM1-3 516 R44 (DM),* and was initially published in 2011^4^ and continuously improved over the past years including a recent update based on long read sequencing^5,6^. Another major advancement in potato genomics was the recent assembly of a heterozygous diploid potato, *RH89-039-16 (RH)*^7^. This haplotype-resolved genome was generated from a variety of different sequencing technologies and phase information from a genetic map derived from selfed progeny.

However, as of now, there is no haplotype-resolved assembly of a tetraploid potato cultivar available nor there is a straightforward method that would enable the assembly of the individual haplotypes of a tetraploid genome. The latest methods for haplotype phasing include the separation of sequencing reads based on the differences between the parental genomes^8^ or based on haplotype information derived from gamete^9–12^ or offspring genomes^7,13^. Similarly, chromosome conformation capture sequencing (e.g., Hi-C) can help to resolve haplotypes during or before the assembly^14–18^ and has been applied to polyploids already^15–17^. However, even though straightforward in its application, chromosome conformation capture sequencing can lead to haplotype switch errors, and requires additional efforts such genetic maps for correction^7,9,18^.

## Genome assembly of a tetraploid potato

We generated an assembly of the autotetraploid genome of *S. tuberosum* ‘Otava’ using high-quality long PacBio HiFi reads (30x per haplotype) using *hifiasm*^19^ (Fig. 1; Data S1; Fig. S1-2; Methods). The initial assembly consisted of 6,366 contigs with an N50 of 2.1 Mb (Fig. S3). While the total assembly size of 2.2 Gb was much larger than the estimated haploid genome size of ~840 Mb, it accounted only for ~65% of the tetraploid genome size (Fig. S2) indicating that one third of the genome collapsed during the assembly. A sequencing depth histogram across the contigs featured four distinct peaks, which originated from regions with either one, two, three, or four (collapsed) haplotype(s) (Fig. 1b). While most of the contigs represented only one haplotype (referred to as *haplotigs)* and accounted for 1.5 Gb (68%) of the assembly, contigs representing two, three or even four collapsed haplotypes (referred to as *diplotigs, triplotigs* or *tetraplotigs)* still made up 470 Mb (21 %), 173 Mb (8%) or 43 Mb (2%). Regions with even higher coverages were virtually absent (9.4 Mb, 0.4%).

**Figure 1.**
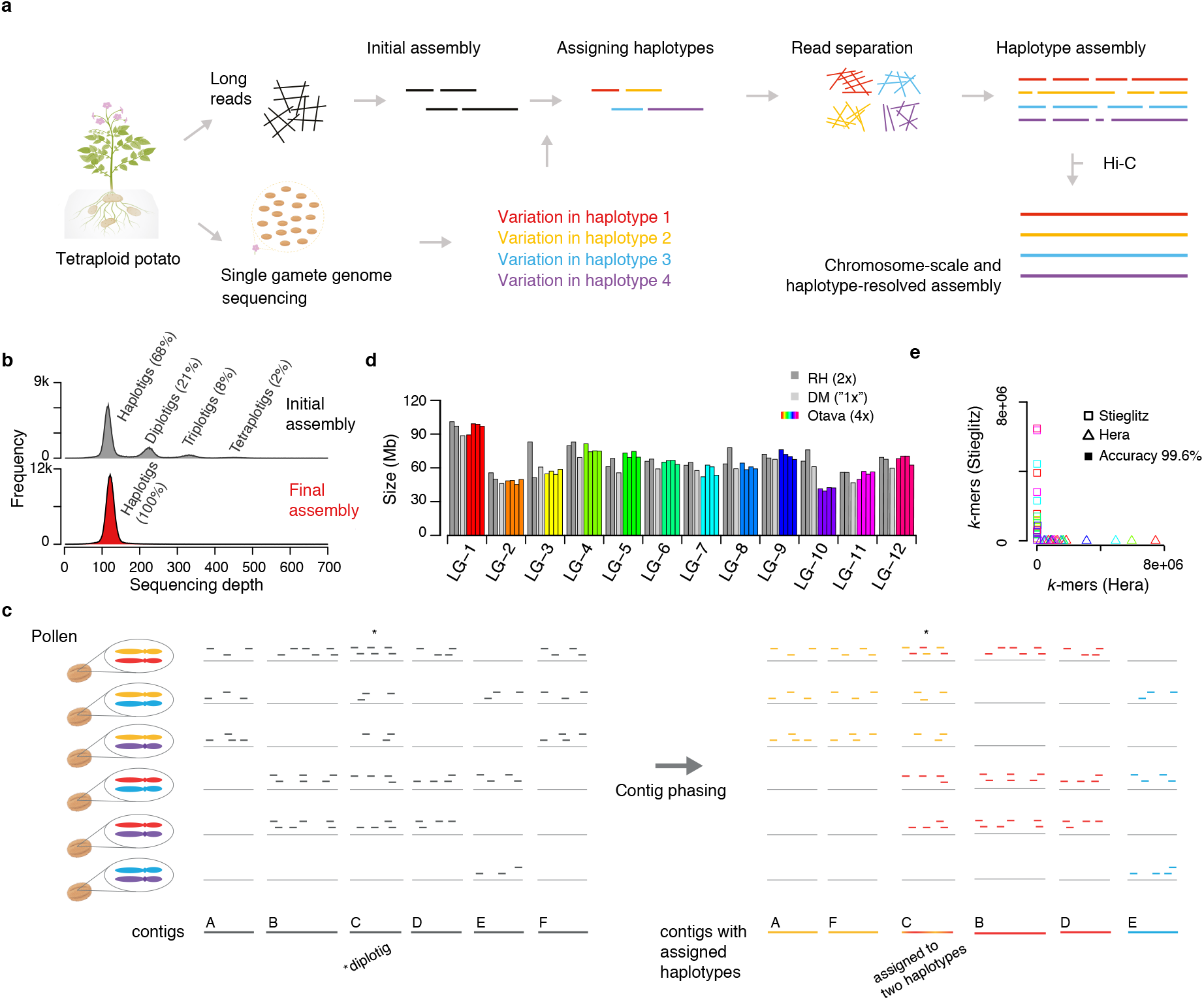
Haplotype-resolved assembly of an autotetraploid potato genome. **a.** Assembly strategy (gamete binning) for tetraploid genomes. Long reads are sequenced from somatic DNA and an initial contig-level assembly is generated. In addition, sequencing data of gamete genomes are generated. Genetic linkage enables grouping of the contig into clusters, which represent the individual haplotypes. Long reads are assigned to haplotypes based on their similarity to the contigs. Each haplotype can be assembled separately and scaffolded to chromosome-scale using Hi-C. (The figure was created with help of BioRender.com.) **b.** Histogram of sequencing depth within 10 kb windows of the initial assembly (top) revealed the presence of haplotigs (68.3%), diplotigs (21.4%), triplotigs (7.9%) and tetraplotigs (2.0%). As a comparison, only one major peak was observed (with increased frequency) in the final assembly (bottom). **c.** Linkage grouping. Presence/absence patterns (PAP) at each coverage marker (50 kb region) are defined by the absence or presence of corresponding sequencing reads from each of the pollen genomes. For instance, the PAP of contig A is “111000”, where “1” refers to pollen genomes with reads that align to the contig and “0” refers to pollen genomes without such reads. PAPs of coverage markers A and F are highly correlated and can be grouped as {A, F}. Similarly, marker B and D are grouped as {B, D}, while coverage marker E remains ungrouped as {E}. The diplotig coverage marker C shows highest correlations to {A, F} and {B, D} as compared to {E}, and therefore extends these two larger clusters. Final result are the three clusters {A, C, F}, {B, C, D} and {E}. **d**. Assembly sizes of the haplotypes, which were highly consistent to the *DM*^5^ and *RH*^7^ assemblies. **e**. *k*-mer based evaluation of the haplotyping accuracy. Each point represents one haplotype of one individual chromosome. The values on the x- and y-axes indicate the numbers of *k*-mers within the haplotype sequence that are unique to either of the parental genomes ‘Hera’ or ‘Stieglitz’. Overall, 99.6% of the variation were correctly phased.

As there is no straight forward solution to untangle collapsed contigs after the assembly, we restarted the genome assembly, but this time based on four separated read sets each derived from one of the four haplotypes. In diploids, such a read separation prior to the assembly can be performed by sorting the reads according to their similarity to the parental genomes (trio binning)^8^. But as autotetraploid individuals inherit two haplotypes through both the maternal and paternal lineages, this cannot be applied for autotetraploid genomes. Alternatively, the reads can also be separated using the haplotypes found in gamete genomes (gamete binning)^9^. While this is straightforward with haploid gametes from diploid individuals, tetraploid potato develops diploid gametes, which again does not separate individual haplotypes. However, as the pairing of the two haplotypes in a diploid gamete is random in potato, we speculated that it might be possible to gain information on individual haplotypes (and thus to separate the reads into four distinct sets) if we sequence a sufficient number of diploid gametes.

To test if gamete binning could be applied for the genome assembly of ‘Otava’, we sequenced the genomes from 717 pollen nuclei with Illumina short reads with an average sequence coverage of 0.18x (Fig. S4) and aligned each of the 717 read sets against the initial assembly. As defining a high-density SNP list can be difficult in a highly heterozygous autotetraploid genome, we defined “coverage markers” (using average alignment depth in 50 kb windows) to assess if a genomic region was present in a pollen genome or not (Methods).

A coverage marker will be covered by reads if one of the two haplotypes of a pollen carries the region of the coverage marker. With this, we could assess the presence/absence pattern (PAP) of a coverage marker across all the 717 pollen genomes (Fig.1c). Closely linked markers featured highly similar PAPs, as most pollen genomes carried the same pair of haplotypes at two neighboring loci. Occasionally, recombination breakpoints, which are integrated into the pollen genomes during meiosis, change the haplotype within a pollen genome, and thereby slightly change the PAPs along the chromosome. However, as recombination is generally rare, closely linked coverage markers still feature highly correlated PAPs. We therefore could use the similarities between the PAPs to cluster the contigs into 48 groups representing the four haplotypes of all 12 chromosomes (Fig. S5-6). Haplotigs were assigned to single clusters. Diplotigs, triplotigs and tetraplotigs represented multiple haplotypes and were therefore assigned to two, three or four of the clusters (Methods).

Once the contigs were assigned to haplotypes, also the PacBio HiFi reads could be assigned to these haplotypes based on their alignments against the contigs. Reads aligned to diplotigs, triplotigs or tetraplotigs were randomly assigned to one of the respective haplotypes. With this, more than 99.9% of the non-organellar PacBio HiFi reads could be assigned to one of the 48 read sets (Fig. S7; Methods). Assembling the read sets using *hifiasm* resulted in 48 haplotype-resolved assemblies with an average N50 of 7.1 Mb and a total size of 3.1 Gb (92% of the tetraploid genome). Finally, we used Hi-C short read data (130x per haplotype) to scaffold the contigs of each assembly to a chromosome-scale, haplotype-resolved assembly (Fig. S8; Methods). Comparison of the full assembly to whole genome sequencing short reads of ‘Otava’ using *Merqury*^20^ revealed very high base accuracy (QV>51.7) and completeness (97.3%) of the ‘Otava’ genome (Methods).

The sizes of the four haplotypes of each chromosome were highly consistent to each other as well as to those of the *DM* and *RH* assemblies^4–7^ except for the consistently shorter assemblies of LG10, which indicated the presence of large-scale chromosomal rearrangements between different cultivars, similar to those previously described^21^ (Fig. 1d). Apart from the LG10 differences, the ‘Otava’ assembly was in high synteny to the DM reference sequence suggesting that also the structure of the chromosomes was assembled correctly (Fig. S9-10). To evaluate the haplotyping accuracy of the tetraploid assembly in more depth, we sequenced the parental cultivars of ‘Otava’, called ‘Stieglitz’ and ‘Hera’, with Illumina short reads at 10x coverage per haplotype, as each of the chromosomes was either inherited from ‘Stieglitz’ or from ‘Hera’. Comparing the *k*-mers, which are specific to one of the parental genomes with each of the 48 chromosome assemblies, we found that each chromosome included almost exclusively *k*-mers from one but not the other parent implying a haplotyping accuracy of 99.6% (Fig. 1e; Methods).

Integrating *ab initio* predictions, protein and RNA-seq read^4–7^ alignments, we annotated 152,855 gene models across four haplotypes, with an overall *BUSCO*^22^ completeness score of 97.3%, which is highly comparable to the annotations of the *RH* and *DM* assemblies^5,7^(Data S2-3; Methods). In addition, we found comparable amounts of various different types of non-coding RNA for all haplotype genomes, which in total accounted for 33.9 Mb across the entire genome (Data S4). Repetitive sequences made up 66% of the assembly with LTR retrotransposons as the most abundant class and rDNA clusters of up to 600 kb in size, which were assembled without any gaps (Data S5-6; Methods). The distribution of genes and repeats along the chromosome followed the typical distribution of mono-centric plant genomes with high gene and low repeat densities at the distal parts of the chromosome, while in the pericentromeric regions the gene densities were low and the repeat densities were high (Fig. 2).

**Figure 2.**
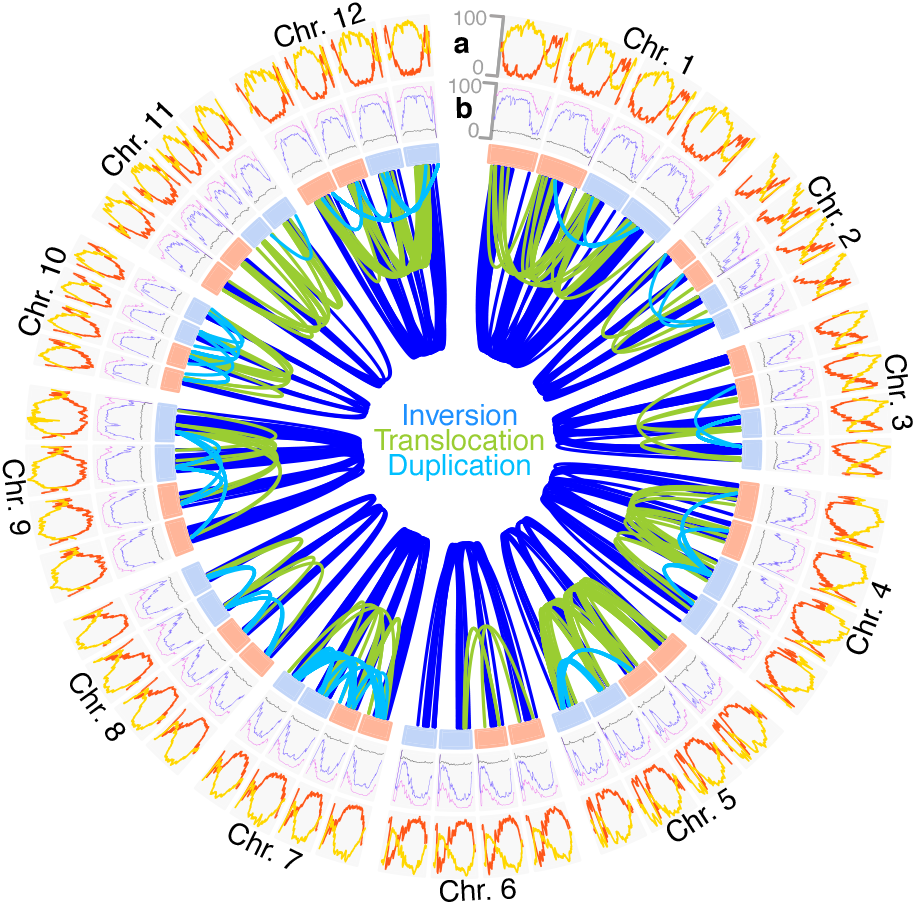
The genomic features of the autotetraploid potato genome. **a**. Gene density (number of genes per Mb; red) and percentage of transposable element-related sequence (yellow) within 2 Mb windows along the four haplotypes of each of the 12 chromosomes. **b**. Landscapes of methylation in CG (purple), CHG (blue), CHH (gray) context within 2 Mb windows. The links in the center show over 600 structural rearrangements larger than 100 kb found between the four haplotypes of each chromosome. Light blue and red box refer to the maternally and paternally inherited chromosomes.

## The genomic footprints of inbreeding

A histogram of sequence differences within 10kb windows between the haplotypes revealed two separated peaks implying the presence of highly similar as well as highly different regions (Fig. 3a). The divergent regions included 1 SNP per 60 bp on average, while the remaining 50% of the regions were almost without any differences (Fig. 3a). This extreme similarity between some of the haplotypes suggested that they were recently inherited from a common ancestor. In fact, the pedigree of many of the cultivated potatoes, including ‘Otava’, contain cultivars that occur more than once in their ancestry^23,24^ (Fig. S1). Common ancestors in different lineages of the pedigree leads to inbreeding and results in regions which are identical-by-descent (IBD) between their haplotypes (Fig. 3b-c; Fig. S11-22; Methods). The IBD blocks almost perfectly matched the collapsed regions in the initial assembly explaining the high degree of unresolved regions. To exclude any potential artefacts, we screened the IBD blocks for collapsed variation (using pseudo-heterozygous variation in read alignments against the assembly) and found that a maximum of 1 SNP in 72 kb could have been missed (Data S7), reassuring that highly similar IBD regions do exist in the genome.

**Figure 3.**
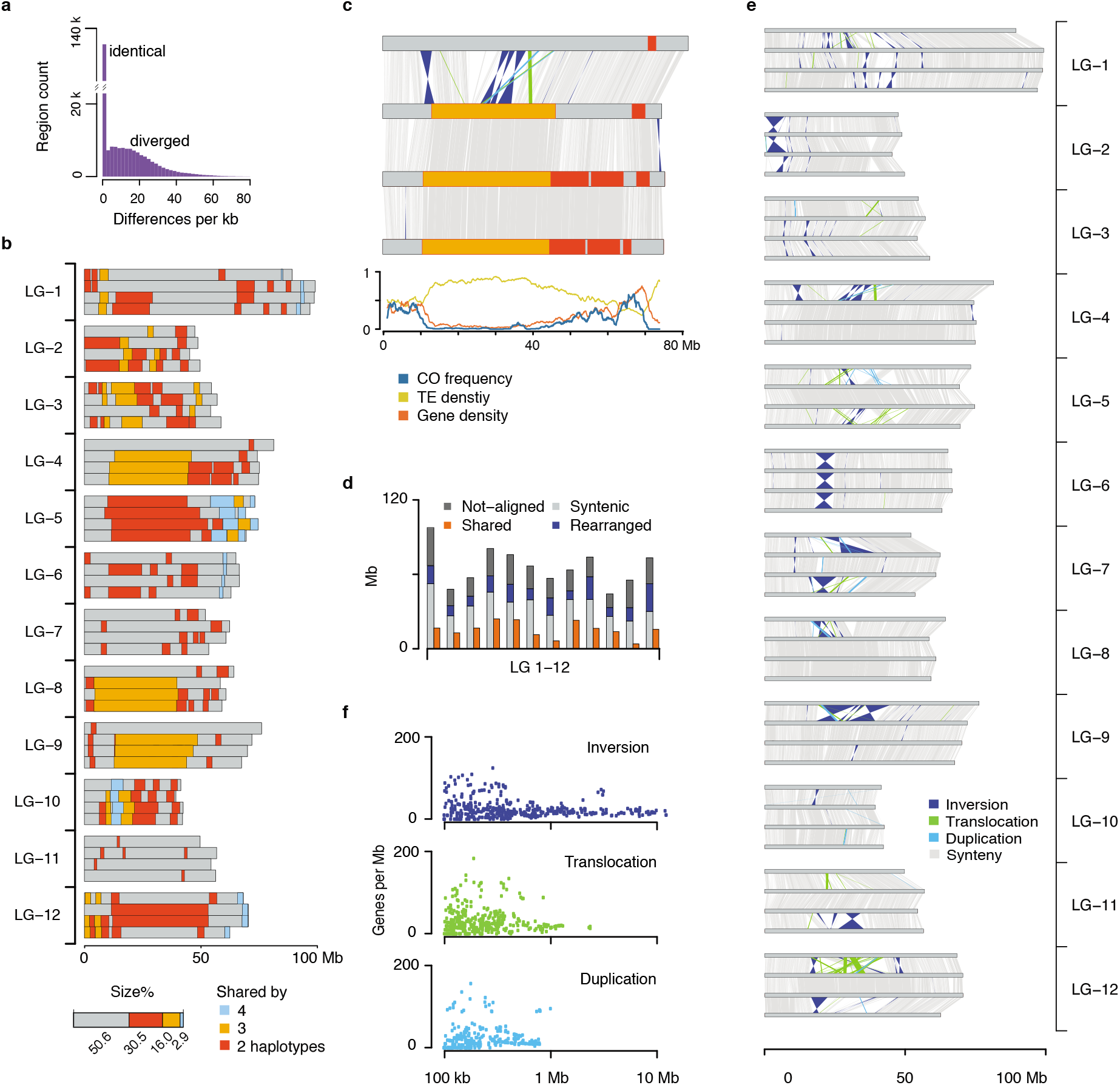
In-depth haplotype analysis of the tetraploid genome. **a.** SNP density as observed in pairwise comparisons between the haplotypes revealed two separated peaks. The high abundance of highly similar/identical regions suggested the existence of identical-by-descent (IBD) blocks. **b.** IBD blocks (minimum size: 1 Mb) across the genome. Regions shared by two, three or four haplotypes are colored in red, orange or blue. **c.** A zoom-in on the IBD blocks and structural rearrangements of LG-4. Large IBD blocks were more likely to occur in peri-centromeric regions with low gene, but high TE content and suppressed meiotic recombination. (Colors as defined in **b** and **e**) **d.** Average alignment statistics and structural rearrangements in each chromosome. **e.** Structural rearrangements between the four haplotypes of each chromosome. **f.** Correlation of the individual size of 220 duplications, 207 translocations and 234 inversions with the respective gene density.

Overall, almost 50% of the tetraploid genome of ‘Otava’ were included in IBD blocks and were shared by either two, three or in rare cases even by four haplotypes (Fig. 3b-d). Individual IBD blocks varied in size and reached up to 41.6 Mb, while IBD blocks in the pericentromeres were significantly larger as compared to the IBD blocks in the distal parts of the chromosomes (Fig. S11-23). Even though it is possible that long IDB blocks were recently introduced and were not broken up by meiotic recombination yet, it is more likely that these extremely long IBD blocks exist due to local suppression of meiotic recombination in the pericentromeres (Fig. 3c). Using the accumulated mutation rates in the IBD blocks as an estimate of their age showed that long IBD blocks weren’t younger as compared to short IBD blocks (Fig. S23b).

## Extreme haplotype differences and their influence on genes

The highly similar IBD blocks were contrasted by high levels of structural rearrangements between the non-shared regions of the genome (Fig. 3; Fig. S11-22; Methods). Inversions, duplications, and translocations made up 3.8% to 42.9% of each of the haplotypes (or 19.3% of the genome) depending on the abundance of IDB blocks in the respective haplotypes. Duplications and translocations were highly enriched for Gypsy and Copia retrotransposons near their breakpoints suggesting their dynamic contribution to genome diversification as it was described for other plant genomes before^25^, while inversions were not enriched for transposable elements (Fig. 3d; Fig. S24; Data S8).

Excluding IBD blocks, structural rearrangements made up 15.0% to 65.8% of each chromosome. In addition, each haplotype included 11.0% to 42.5% of unique sequence that could not be aligned to any of the other haplotypes (Fig. 3d). This amount of structural variation and haplotype-specific sequence was much higher than what has been reported for any other crop species so far, supporting earlier suggestions that genomic introgressions from wild relatives were part of the domestication history of potato^26^.

Overall, we found 661 structural variations longer than 100 kb which all were supported by the contiguity of the assembled contigs or Hi-C contact signals, including 220 duplications, 207 translocations and 234 inversions (Fig. 2; Fig. S8,11-22,25; Data S9). While comparable in number, inversions were much larger than the other types of rearrangements and reached sizes of up to 12.4 Mb (Fig. 3e-f). Although these large inversions were mostly located in the peri-centromeric regions where genes occur at low density, they still harbored nearly 5% of all genes (7,958 out of 152,855). Meiotic crossover events from the pollen genomes were virtually absent in inversions, indicating that these regions are likely to introduce large segregating haplotypes among cultivated potato (Fig. 3c).

Pairwise allelic divergence of the genes ranged from 0 to 140 differences per kb and included identical as well as divergent alleles. The average pairwise difference of the divergent alleles was 18 differences per kb (Fig. 4a) and only 53.6% of the genes were present in all four haplotypes. The remaining 46.4% of the genes were present in three (20.0%), two (15.9%) or even only one (10.5%) of the haplotypes (Fig. 4b) with an average of 3.2 copies per gene. In addition, the coding sequences of some of these copies were identical to each other. For example, only 3,066 (15.4%) of the genes with four copies also featured four distinct alleles. In consequence, even though each gene featured 3.2 copies on average, there were only 1.9 distinct alleles per gene (Fig. 4b).

**Figure 4.**
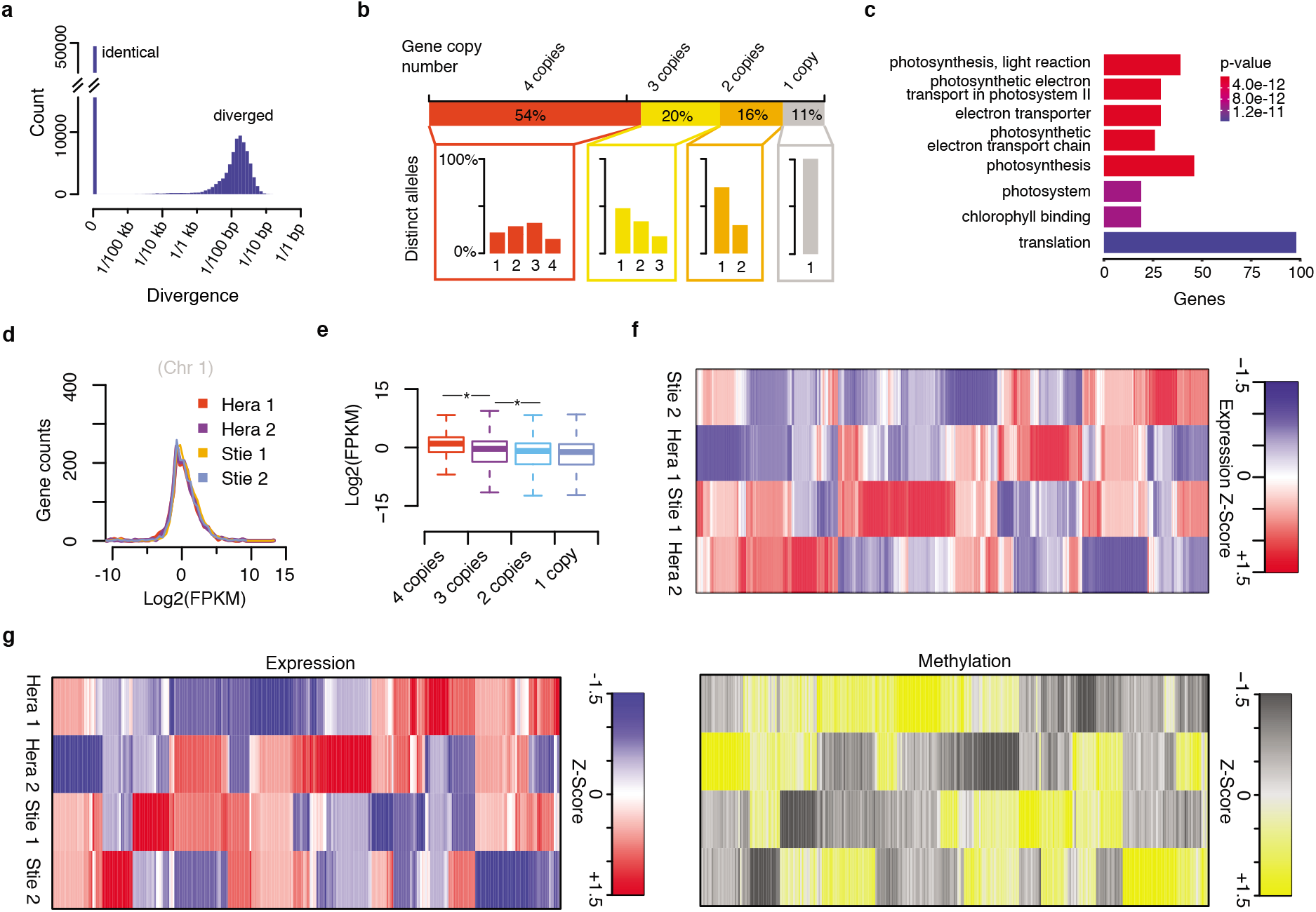
Impact of haplotype divergence on genes and their expression. **a.** Pair-wise evaluation of allelic divergence of genes. **b.** Presence/absence variations of genes. Overall, 53.6%, 20.0%, 15.9% or 10.5% of the genes showed four, three, two or one allelic copies/copy within the tetraploid genome with an average of 3.2 allelic copies per gene. Different configurations of divergent alleles were observed for the sets of genes with four, three or two copies, for instance, within the genes with four allelic copies, one (22.3%), two (29.8%), three (32.5%) or four (15.4%) divergent allele/s were observed. Overall, this led to an average of 1.9 distinct alleles per gene. **c.** GO enrichment analysis of genes with four identical alleles. **d**. Four haplotypes of chromosome 1 showed comparable amounts of expressed transcripts (FPKM: fragments per kilobase per million reads; for chromosome 2-12 see Fig. S26). **e**. Genes with more allelic copies are significantly higher expressed than those with fewer copies (‘*’: *t*-test, *p-* value<0.01). **f**. Among the four haplotypes, 10.8% (1,219) of the 11,154 genes with four functional alleles showed allele-specific expression (Data S11). **g**. Among those, allele-specific expression of 304 genes was correlated with DNA methylation levels, which were measured in the 1 kb up- or downstream regions surrounding the genes (Data S12).

While it was expected to find identical gene alleles within the IBD blocks, only ~45% of the identical gene alleles were actually within shared regions. To test if the high number of identical alleles between the otherwise different haplotypes was indicative of selection, we tested whether these genes were enriched for specific functions. This revealed a significant enrichment for genes with GO terms involving photosynthesis, chlorophyll binding and translation (Fig. 4c) suggesting a selection-induced reduction of allelic diversity through the optimization of plant performance.

The low number of distinct alleles per gene and a selection-induced reduction of allelic difference also implied that the tetraploid nature of the genome is not a necessary feature of the high performance of potato in different environments. However, transforming potato into a diploid suffers from the random distribution of the non-functional alleles throughout the individual haplotypes implying that any ploidy reduction would lead to a significant gene loss. In fact, the *BUSCO* score (indicating completeness) of the annotations of the individual haplotypes was 89.5% on average, while the score of all four haplotypes combined was 97.3% (Data S3) evidencing that the individual haplotypes lack genes which are present elsewhere in the genome. Likewise, the doubled-monoploid *DM*^4,5^ and the diploid *RH*^7^ genomes, which both were derived from tetraploid cultivars, feature 5,901 (15.4%) or 3,245 (8.5%), respectively, less genes as compared to the tetraploid genome. The gene family with highest percentage of genes with presence/absence variation (45.4%; 316 out of 696 genes) were the NLR resistance genes (Data S10), which are known for their high intraspecies variability^27,28^.

To investigate how genes are expressed in this tetraploid genome, we sequenced 367 million read pairs of the ‘Otava’ leaf transcriptome in three replicates (Data S1; Methods). The four haplotype genomes contributed highly similar amounts of RNA, suggesting that none of the haplotypes was dominant (Fig. 4d; Fig. S26), which is similar to observations in another autoploid species^15^. Comparable to earlier analyses on the effects of CNVs on gene expression^29^, the number of allelic copies also impacted on gene expression. Genes with more allelic copies showed a significantly increased gene expression as compared to genes with fewer allelic copies (Fig. 4e). Although gene expression of the four haplotype genomes were comparable at genome scale, 10.9% of the genes with four allelic copies featured significant expression differences between the individual alleles (Fig. 4f).

To understand more about the regulation of allele-specific expression, we sequenced the DNA methylome of ‘Otava’ using enzymatic methylome sequencing with three replicates, each with 277 million read pairs (Data S1; Methods). Overall, we found that DNA methylation was consistent across all haplotypes while DNA methylation levels in IBD blocks were slightly higher as compared to the non-shared regions in the other haplotypes (Fig. 2; Fig. S27). Of the 1,219 genes with significant differences in the allele-specific expression, 304 genes were significantly correlated with the level of methylation at the up/downstream regions of these genes (Fig. 4g), suggesting that around 25% of the allele-specific expression are regulated through DNA methylation.

## Discussions

Here we reported the first haplotype-resolved assembly of an autotetraploid potato. Leveraging high-quality, long reads and single-cell genotyping of diploid gametes, we were able to reconstruct the sequences of all four haplotypes. The sequence differences between the haplotypes were much higher as compared to the differences commonly found within species and were rather reminiscent of the differences found between species. This supports earlier suggestions that the diverse haplotypes originated from introgressions from divergent species during domestication^26^.

The high level of sequence differences, however, was contrasted by widespread IBD blocks, which were most likely introduced by related genotypes during breeding, even though we cannot exclude that some of these blocks might have been formed via double reduction during meiosis^30^. This similarity of the IBD blocks was the reason for the abundant collapsed regions in the initial assembly. As these regions were almost identical, it was not possible to assemble them from the sequence data alone. IBD blocks are a widespread phenomenon in many crops or livestock in general, though the challenges associated with the high similarity between haplotypes can be solved by using the power of genetics and analyzing individual gamete genomes.

The abundance of IBD blocks, in addition to IBD-independent allele sharing, led to the unexpected observation that the tetraploid genome only included 1.9 diverse allelic copies per gene. This implied that the maximal allelic diversity that could be included in the tetraploid genome was not reached, even though the high yield and yield stability of potato is supposed to be promoted by the effects of heterosis, which itself is based on non-additive interactions of diverse alleles^31^. Whether the high abundance of shared alleles suggests that the effects of heterosis could still be optimized by increasing the number of polymorphic alleles or if this indicates that the limits of heterosis were already reached remains to be seen.

Over the past years, considerable success has been made in re-domesticating potato from a clonally-propagated, tetraploid crop into a seed-propagated, diploid crop to increase reproduction rate, decrease costs in storage and transportation, and improve disease control^2,32–34^. However, the random distribution of loss-of-function alleles in tetraploid potato can lead to the accelerated manifestation of inbreeding depression in the diploid genomes, when they are derived from tetraploids^7,35^. Haplotype-resolved assemblies of autotetraploids like the one presented here have the potential to support the design of optimal haplotypes by avoiding the combination of known incompatibility alleles^36^. Of course, this new possibility to assemble autotetraploid genomes does not eliminate all breeding-related problems that result from the tetraploid nature of potato. However, being able to reconstruct the four haplotypes of cultivated potato is a breakthrough for modern genomics-assisted breeding strategies, and ultimately has the power to increase the breeding success of potato in the future.

## Methods

Plant material was grown at Max Planck Institute for Plant Breeding Research (Cologne, Germany). The genome of ‘Otava’ was sequenced with PacBio HiFi Sequel II platform with four SMRTcells. DNA extracted from individual pollen nuclei was prepared with 10x Genomics CNV kits and subsequently sequenced with Illumina sequencing. Barcodes (in single cell sequencing) were corrected using *cellranger* (10x Genomics). Short/long reads were aligned using *bowtie2*^37^ *Iminimap2*^38^. BAM, VCF file processing and sequencing depth analysis were performed using *samtools^39^* and *bedtools*^40^. Linkage grouping was performed using in-house developed code. PacBio sequence reads were assembled using *hifiasm*^19^, and genome annotation was performed following a previous pipeline^9^. Structural variations were identified using *SyRI*^41^ based on *minimap2* genome alignments. Expression analysis was performed following the literature^42^. Methylation calling was performed with *bismark* pipeline^43^. More details and other related methods are provided in the Supplementary Information.

## Acknowledgements

The authors would like to thank Christiane Gebhardt (MPI-PZ, Cologne, Germany) and Benjamin Stich (HHU, Düsseldorf, Germany) for helpful discussions, Birgit Walkemeier and Christine Sänger (both MPI-PZ, Cologne, Germany) for plant cultivation, Christine Brandt and Klaus J. Dehmer (both IPK, Groß Lüsewitz, Germany) for providing material, Pádraic J. Flood (WUR, Wageningen, The Netherlands) for comments on the manuscript as well as Saurabh Pophaly (MPI-PZ, Cologne, Germany) for enthusiastic help in data management. We are very grateful to John Hamilton and C. Robin Buell (both University of Georgia, Georgia, USA) for integrating the ‘Otava’ assembly into Spud DB^6^.

## Author contributions

H.S. and K.S. developed the project. J.A.C., H.S. and K.S. coordinated the generation of data with K.K., K.F-D., C.K. and B.H.. H.S., W-B.J., and M.G. performed all data analysis. H.S. and K.S. wrote the manuscript with input from all authors. All authors read and approved the final manuscript.

## Data and materials availability

High-throughput sequencing data analyzed in this project are available under NCBI BioProject: PRJNA751899. This Whole Genome Shotgun project (including assembly and annotation) has been deposited at DDBJ/ENA/GenBank under the accessions JAIVGA000000000, JAIVGB000000000, JAIVGC000000000 and JAIVGD000000000. The version described in this paper is version JAIVGA010000000, JAIVGB010000000, JAIVGC010000000 and JAIVGD010000000. The genome assembly and gene annotation of ‘Otava’ are also available on Spud DB (http://spuddb.uga.edu/)^6^. Customed code and scripts supporting this work are available at github.com/schneeberger-lab/GameteBinning_tetraploid.

## Supplementary Information

Materials and Methods

Figs. S1 to S28

Data S1 to S12

## References

1. The Food and Agriculture Organization (FAO). http://www.fao.org/faostat/en/#data/QV (2021).

2. S. H. Jansky et al. Reinventing potato as a diploid inbred line-based crop. Crop Sci. 56, 1412–1422 (2016).

3. D.S. Douches, D. Maas, K. Jastrzebski, R.W. Chase. Assessment of Potato Breeding Progress in the USA over the Last Century. Crop Sci. 36, 1544–1552 (1996).

4. The Potato Genome Sequencing Consortium. Genome sequence and analysis of the tuber crop potato. Nature 475, 189–195 (2011).

5. G.M. Pham, J.P. Hamilton et al. Construction of a chromosome-scale long-read reference genome assembly for potato. GigaScience 9, 1–11 (2020).

6. C.D. Hirsch, J.P. Hamilton, K.L. Childs, J. Cepela, E. Crisovan, et al. (2014), Spud DB: A Resource for Mining Sequences, Genotypes, and Phenotypes to Accelerate Potato Breeding. The Plant Genome 7, plantgenome2013.12.0042 (2014).

7. Q. Zhou, D. Tang et al. Haplotype-resolved genome analyses of a heterozygous diploid potato. Nat. Genet. 52, 1018–1023 (2020).

8. S. Koren, A. Rhie et al. De novo assembly of haplotype-resolved genomes with trio binning. Nat. Biotechnol. 36, 1174–1182 (2018).

9. J.A. Campoy, H. Sun et al. *Gamete binning:* chromosome-level and haplotype-resolved genome assembly enabled by high-throughput single-cell sequencing of gamete genomes. Genome Biol. 21, 306 (2020).

10. R. Li, H. Qu et al. Inference of chromosome-length haplotypes using genomic data of three or a few more single gametes. Mol Biol Evol. 37, 3684–3698 (2020).

11. E.F. Kirkness et al. Sequencing of isolated sperm cells for direct haplotyping of a human genome. Genome Res. 23, 826–832 (2013).

12. D. Shi, J. Wu, H. Tang, H. Yin et al. Single-pollen-cell sequencing for gamete-based phased diploid genome assembly in plants. Genome Res. 29, 1889–1899 (2019).

13. C. Zhou, et al. Assembly of whole-chromosome pseudomolecules for polyploid plant genomes using outbred mapping populations. Nat. Genet. 52, 1256–1264 (2020).

14. S. Garg, et al. Chromosome-scale, haplotype-resolved assembly of human genomes. Nat. Biotechnol. (2020).

15. J. Zhang, X. Zhang, H. Tang, Q. Zhang et al. Allele-defined genome of the autopolyploid sugarcane *Saccharum spontaneum* L. Nat. Genet. 50, 1565–1573 (2018).

16. H. Chen, Y. Zeng, Y. Yang, L. Huang, B. Tang, H. Zhang. et al. Allele-aware chromosome-level genome assembly and efficient transgene-free genome editing for the autotetraploid cultivated alfalfa. Nat Commun 11, 2494 (2020).

17. X. Zhang, S. Zhang, Q. Zhao, R. Ming, H. Tang. Assembly of allele-aware, chromosomal-scale autopolyploid genomes based on Hi-C data. Nat. Plants 5, 833–845 (2019).

18. G. Linsmith et al. Pseudo-chromosome-length genome assembly of a double haploid *“Bartlett”* pear *(Pyrus communis L.)*. GigaScience 8, 1–17 (2019).

19. H. Cheng, G.T. Concepcion, X. Feng, H. Zhang, H. Li. Haplotype-resolved de novo assembly with phased assembly graphs. Nat Methods 18, 170–175 (2021).

20. A. Rhie, B.P. Walenz, S. Koren, A.M. Phillippy. *Merqury:* reference-free quality, completeness, and phasing assessment for genome assemblies. Genome Biol 21, 1–27 (2020).

21. L. Comai, K. Amundson, B. Ordonez, X. Zhao, G. Tomaz Braz, J. Jiang, I. Henry. LD-CNV: rapid and simple discovery of chromosomal translocations using linkage disequilibrium between copy number variable loci. Preprint at Biorxiv (2021).

22. F.A. Simão and R.M. Waterhouse et al. *BUSCO:* Assessing genome assembly and annotation completeness with single-copy orthologs. Bioinformatics 31, 3210–3212 (2015).

23. R.C.B. Hutten and R. van Berloo. An online potato pedigree database. URL: http://www.plantbreeding.wur.nl/PotatoPedigree/ (2001).

24. R. van Berloo, R.C.B. Hutten, H.J. van Eck, and R.G.F Visser. An online potato pedigree database resource. Potato research 50, 45–57 (2007).

25. E.M. Willing, V. Rawat, T. Mandáková et al. Genome expansion of *Arabis alpina* linked with retrotransposition and reduced symmetric DNA methylation. Nature Plants 1, 14023 (2015).

26. Ma.A. Hardigan, F.P.E. Laimbeer, L. Newton, E. Crisovan, J.P. Hamilton, B. Vaillancourt et al. Genome diversity of tuber-bearing Solanum uncovers complex evolutionary history and targets of domestication in the cultivated potato. Proc Natl Acad Sci U S A 114, E9999–E10008 (2017). doi: 10.1073/pnas.1714380114.

27. A.L. Van de Weyer, F. Monteiro, O.J. Furzer, et al. A Species-Wide Inventory of NLR Genes and Alleles in Arabidopsis thaliana. Cell. 178, 1260–1272 (2019).

28. K. Seong, E. Seo, K. Witek, M. Li, B. Staskawicz. Evolution of NLR resistance genes with noncanonical N-terminal domains in wild tomato species. New Phytol. 227, 1530–1543 (2020).

29. G.M. Pham, L. Newton, K. Wiegert-Rininger, B. Vaillancourt, D.S. Douches, C.R. Buell. Extensive genome heterogeneity leads to preferential allele expression and copy numberdependent expression in cultivated potato. Plant J. 92, 624–637 (2017).

30. P.M. Bourke, R.E. Voorrips, R.G. Visser, C. Maliepaard. The double-reduction landscape in tetraploid potato as revealed by a high-density linkage map. Genetics, 201:853–863 (2015).

31. J. Muthoni, H. Shimelis, R. Melis. Production of hybrid potatoes: Are heterozygosity and ploidy levels important? Australian Journal of Crop Science 13, 687–694 (2019).

32. P. Lindhout. et al. Towards F1 Hybrid Seed Potato Breeding. Potato Res. 54, 301–312 (2011).

33. M. Ye, Z. Peng et al. Generation of self-compatible diploid potato by knockout of S-RNase. Nature Plants 4, 651–654 (2018).

34. Y. Li, G. Li, C. Li, D. Qu., S. Huang. Prospects of diploid hybrid breeding in potato. Chin. Potato J. 27, 96–99 (2013).

35. C. Zhang, P. Wang, D. Tang et al. The genetic basis of inbreeding depression in potato. Nat Genet. 51, 374–378 (2019).

36. Q. Lian, D. Tang, Z. Bai, J. Qi, F. Lu, S. Huang, C. Zhang. Acquisition of deleterious mutations during potato polyploidization. J Integr Plant Biol. 61, 7–11 (2019).

37. B. Langmead, S.L. Salzberg. Fast gapped-read alignment with *Bowtie* 2. Nature methods 9, 357–359 (2012).

38. H. Li. *Minimap2:* Pairwise alignment for nucleotide sequences. Bioinformatics 34(18), 3094–100 (2018).

39. H. Li. et al. The Sequence Alignment/Map format and SAMtools. Bioinformatics 25, 2078–2079 (2009).

40. A.R. Quinlan, I.M. Hall. *BEDTools:* A flexible suite of utilities for comparing genomic features. Bioinformatics 26, 841–842 (2010).

41. M. Goel, H. Sun, W.B. Jiao, K. Schneeberger. *SyRI*: finding genomic rearrangements and local sequence differences from whole-genome assemblies. Genome Biol. 20, 1–13 (2019).

42. C.W. Law, M. Alhamdoosh, S. Su, X. Dong, L. Tian et al. RNA-seq analysis is easy as 1-2-3 with limma, Glimma and edgeR. F1000Research 5, ISCB Comm J-1408 (2016).

43. F. Krueger, S.R. Andrews. *Bismark*: a flexible aligner and methylation caller for Bisulfite-Seq applications. Bioinformatics 27, 1571–2 (2011).

