## Supplementary Information for "Chromosome-scale and haplotype-resolved genome assembly of a tetraploid potato cultivar"

**This file includes:**

Materials and Methods

Figure S1 to 28

**Other Supplementary Materials for this manuscript include the following:**

Data S1-12

### Materials and Methods

#### Plant Material

The three potato cultivars ‘Otava’, ‘Hera’ and ‘Stieglitz’ (Figure S1) were clonally propagated and grown on Murashige-Skoog-Medium (MS) for 3-4 weeks at Max Planck Institute for Plant Breeding Research (MPIPZ, Germany). Seedlings were transferred to soil in 7 x 7 cm pots and grown in a Percival growth chamber for 2-3 weeks. Afterwards, potato plants were transferred to 1-liter pots and grown until flowering. Potatos were grown in long day (LD) conditions (16 h light, 8 h night cycle), at 22 °C.

#### HMW somatic DNA extraction, library preparation and sequencing (‘Otava’)

High-molecular-weight DNA was isolated from 1.5 gram material (Otava leaves) with a NucleoBond HMW DNA kit (Macherey Nagel). Quality was assessed with a FEMTOpulse device (Agilent) and quantity measured by fluorometry Quantus (Promega). A HiFi library was then prepared according to the manual "Procedure & Checklist - Preparing HiFi SMRTbell® Libraries using SMRTbell Express Template Prep Kit 2.0" with initial DNA fragmentation by g-Tubes (Covaris) and final library size binning by SageELF (Sage Science). Size distribution was again controlled by FEMTOpulse (Agilent). Size-selected libraries were sequenced on a Sequel II device at Max Planck Genome-centre Cologne (MP-GC) with Binding kit 2.0 and Sequel II Sequencing Kit 2.0 for 30 h. Details of read information are provided in Data S1.

#### Pollen DNA extraction, library preparation and sequencing (‘Otava’)

Pollen grains of ‘Otava’ were extracted following recent work[1]. Ten anthers from two ‘Otava’ flowers were extracted with forceps and submerged in woody pollen buffer (WPB). Around 800,000 pollen grains were extracted from the anthers by vortexing them in WPB. Isolated pollen was prefiltered (100 µm) and bursted (10 µm) using Celltrics™ sieves and WPB. After sorting and counting using the BD FACSAria™ Fusion flow cytometer with high-speed sort settings (70 µm nozzle and 70 PSI sheath pressure, at Max Planck Institute for Biology of Ageing), a total of 12,600 nuclei were selected and collected in a solution of 4.2 µL phosphate-buffered saline with 0.1% bovine serum albumin. According to manufacturer’s instructions, the nuclei were loaded into a 10x Genomics Chromium Controller in two batches each with 6,300 nuclei. This led to two 10x single cell CNV libraries (DNA), and both were sequenced with a Illumina HiSeq3000 device in 150 bp paired-end read mode at Max Planck Genome center (MPGC, Cologne, Germany), reaching a coverage of 52x per haplotype. Details of read information are provided in Data S1.

#### Omni-C DNA extraction, library preparation and sequencing ('Otava')

An aliquot of HMW DNA was extracted from fresh leaves of 'Otava' used for a Dovetail Omni-C library created at MPGC using the Omni-C™ Kit. The library was sent to BGI, Hongkong (China) with dry ice, where it got sequenced on DNBSEQ-G400 platform. Details of read information are provided in Data S1.

#### RNA library preparation and sequencing ('Otava')

RNA was isolated from leaves with an RNeasy plant kit, Qiagen including an on-column RNase treatment. Poly-A RNA was enriched from 1 µg total RNA by the NEBNext® Poly(A) mRNA Magnetic Isolation Module. RNAseq libraries were prepared as described in NEBNext Ultra™ II Directional RNA Library Prep Kit for Illumina (New England Biolabs). A total of eleven cycles were applied to enrich library concentration. Sequencing-by-synthesis was performed on a NextSeq 2000 with P3 chemistry and 2 x 150 bp read mode. Three replicates from three leaves of the same 'Otava' plant were made. Details of read information are provided in Data S1.

#### Enzymatic methylome library preparation and sequencing ('Otava')

For Enzymatic Methyl-seq (EM-seq) genomic DNA was isolated with DNAeasy plant mini kit, Qiagen. Then 200 ng DNA was fragmented with COVARIS S2 to 300 bp including spke-ins as recommended by NEB. An Illumina-compatible library was prepared according to the NEBNext® Enzymatic Methyl-seq Kit protocol (NEB) with a total of 5 cycles to enrich barcoded library fragments. Sequencing-by-synthesis was performed on a NextSeq 2000 with P3 chemistry and 2 x 150 bp read mode. Three replicates from three leaves of the same 'Otava' plant were made. Note, the three leaves here were the same as those used for RNA extraction, where a leaf was cut into 2 pieces with one half for RNA sequencing and the other for methylation sequencing. Details of read information are provided in Data S1.

#### DNA extraction, library preparation for linked-read sequencing

One 10x Genomics linked-read library (DNA load: 0.625 ng) was created using DNA extracted from leaves (NucleoBond HMW DNA kit, Macherey Nagel), after size-selection for ≥45 kb with a Sage Science BluePippin high-pass protocol (U1 marker, 0,75% cassette), respectively for 'Otava', 'Hera' and 'Stieglitz'. The libraries were sequenced on Illumina HiSeq3000 platform in 150 bp paired-end read mode (at MPGC). The 'Otava' sequencing was used in sequencing depth analysis, while the parental sequencings were used in *k*-mer based haplotyping evaluation analysis. Details of read information are provided in Data S1.

#### DNA extraction, library preparation and sequencing (‘Stieglitz’ and ‘Hera’)

Fresh leaves were sampled from the parental cultivar ‘Hera’, DNA was extracted using the Plant DNA Kit of Macherey-Nagel™, treated with RNAse, and an Illumina-compatible short read library was prepared after gDNA fragmentation (S2, Covaris) with an Ovation ultralow V2 library kit (Tecan Genomics). The library was sequenced using Illumina HiSeq3000 (at MPGC) in 150 bp paired-end read mode. Similarly, an Illumina library was prepared and sequenced on the same device for the *Stieglitz* genome (NEBNext® Ultra™ II FS DNA Library Prep Kit for Illumina®). Details of read information are provided in Data S1. Note that all parental genome data were used only for evaluating haplotyping accuracy in the haplotype-specific assemblies.

#### Genome size estimation

After trimming off 10x Genomics barcodes and hexamers from the 370.3 Gb reads combined from the 10x single cell CNV libraries and single-molecule libraries, *k*-mer counting (*k*=21) was performed with *Jellyfish* (version 2.2.10)[2]. The *k*-mer histogram was provided to *findGSE* (version 1.0)[3] to estimate the haploid/tetraploid genome size of ‘Otava’ under the heterozygous mode (with ‘*exp_hom*=200’; Figure S2).

#### Initial tetraploid genome assembly, polishing and purging

The initial assembly of the tetraploid genome was performed using *Hifiasm* (version 0.7)[4] with default settings with the 102.2 Gb raw PacBio HiFi reads of ‘Otava’, where the output consisting of unitigs (i.e., locally haplotype-resolved contigs) was selected for further processing. Then the short reads from the two 10x single cell CNV libraries were aligned to the assembly using *bowtie2* (version 2.2.8)[5]. These alignments were used to polish the assembly with *pilon* (version 1.22)[6] with options of --*fix bases* --*changes* --*diploid* --*mindepth* 0.8. Further, short reads of the two 10x single cell CNV libraries, additional 10x single molecule libraries and PacBio HiFi reads were all aligned to the polished assembly (using *bowtie2* and *minimap2* (version 2.17-r491)[7] respectively). Among 17,153 raw contigs, 9041 contigs which were longer than 50 kb and with an average sequencing depth over 80x were kept. HiFi reads were re-aligned to the purged assembly and contigs covered less than 3x were removed. The HiFi reads-based purging process was repeated for five rounds to get an initial assembly of 6,366 contigs for subsequent analysis (Figure S3).

#### Coverage marker definition by sequencing depth analysis

All Illumina paired-end short reads from 10x single-cell/molecule libraries (and PacBio HiFi long reads) were aligned to the initial assembly of 6,366 contigs respectively using *bowtie2* and *minimap2*. Potentially duplicated short reads were removed using *picard* “*MarkDuplicates*” function (*http://broadinstitute.github.io/picard/*). The depth along each contig was calculated with *samtools depth* function (version 1.9)[8] for each type of (short and long) reads and the depth at each position of each contig was taken as the sum of all types. Each contig was dissected into 10 kb windows and the average sequencing depth per base was calculated within each window. According to the average genome-wide depth of 113x per haplotype (denoted by *H*), windows with depths in [0, 170x], [171x, 283x], [284x, 396x], [397x, 509x] and [510x, INF] were respectively determined as contig types of haplotig, diplotig, triplotig, tetraplotig and replotig, where the upper bound on depth for each type (if any) was determined by $H*(i+1/2)$, with *i* being 1, 2, 3, 4. Neighboring 10 kb windows (along the same contig) were further merged as larger 50 kb coverage markers if they were classified as the same contig type (to ensure sufficient read signal for genotyping within each single pollen genome).

#### 10x Genomics sequencing barcode correction

The improved version of the *DM* potato reference genome v4.04[9,10] was indexed with *cellranger-dna* (version 1.1.0, 10x Genomics), using sub-function *mkref* under default settings. For each of the two 10x single cell CNV libraries, reads were aligned to the *DM* genome using *cellranger-dna cnv* under default settings. The generated bam files were sorted with *samtools* with *-n* option, and reads updated with corrected barcodes were respectively extracted using customized code.

#### Selection of single-cell sequencings

The read sets of each of the single pollen genomes were aligned to the initial assembly of 6,366 contigs using *bowtie2*. After filtering out non-primary reads using *samtools view* with options "*-F* 3840 *-q* 1", read counts within each 50 kb coverage marker along each contigs were obtained using *bedtools* (v2.29.0)[11]. The coverage ratio of the assembly by a read set was calculated as the total size of covered coverage markers divided by the total assembly size (where a marker harboring more than $7*(N/10^6)$ reads was considered as covered, with *N* being the number of reads aligned to the assembly). Finally, 717 read sets each with over 70,000 read pairs (equivalent to 0.02x of the haploid genome size) which could cover the assembly with a ratio of 0.55 to 0.72 were selected to perform haplotype-specific contig grouping (Figure S4). Pooled alignments of these short reads to the initial assembly delivered solid coverage (above 17x) across 98.5% of the initial assembly, supporting that the 10x libraries had captured comprehensive parts of the genome.

#### Linkage-based grouping of contigs

The read count (here, denoted by *r*) at each coverage marker of size *W* (in bp) for each of the 717 pollen genomes was firstly normalized using $n_r = r*(10^4/W)*(10^6/N)$, where *N* was the total number of reads aligned to the assembly of 6,366 contigs. Meanwhile, the average read count, $m_r$, for all coverage markers with more than 7*($N$/$10^6$) reads was calculated for each pollen genome. The coverage (genotype) at a marker for each pollen was set to $n_r/m_r$ (which would be rounded/forced as 0, 1 or 2). In general, coverages across 717 pollen read sets at the same window marker were used to build up a PAP: $X=x_1x_2 \ldots x_{717}$, where $x_i$ was in {0, 1}, {0, 1, 2}, {1, 2} or {2}, depending on whether the marker type was haplotig, diplotig, triplotig or tetraplotig. The correlation between two PAPs $X$ and $Y$ was calculated as $cor_{XY}$ = $sum[\ (x_i\text{-}m_x) * (y_i\text{-}m_y)\ ] / [\ sqrt(\ sum(x_i\text{-}m_x)^2\ ) * sqrt(\ sum(y_i\text{-}m_y)^2\ )\ ]$, where $m_x$ and $m_y$ were the respective average values. Initially, any pair of haplotigs/vertices (with sizes ≥100 kb) was connected by an edge, if the highest correlation value between the PAPs of the markers at two ends of the two haplotigs was larger than 0.55. This graph based clustering led to 48 groups representing the 48 haplotypes (Figure S5-6). If the lowest correlation value between any pair of the markers of any two groups was less than -0.25, they could be determined as homologous linkage groups (same chromosome, different haplotypes). With this, the 48 groups were clustered into 12 chromosomes, each with four different haplotypes (Figure S5-6).

Each of the remaining haplotigs $h$ (<100 kb) was integrated into the group with the marker showing the highest correlation with $h$. For a diplotig marker (with PAP $Z$) which represent the collapsed haplotypes $A_1$ and $A_2$, we can expect that $Z \approx X+Y$, where $X$ and $Y$ are PAPs of two markers which closely linked to the diploid marker (Main text Fig. 1c). We can therefore expect $Z\&X \approx X$ and $Z\&Y \approx Y$, where '&' refers to the bit-wise AND-operation. As a result, the correlations of '$Z\&X$ with $X$' and '$Z\&Y$ with $Y$' should give the two highest values. Therefore, the two coverage markers from two of the 48 groups that show the highest correlations to a diplotig marker reveal the two groups that the diploitig will be assigned to. Similarly, triplotig markers can be associated with three groups. If all coverage markers of a contig are tetraplotig-type, the contig cannot be associated with any group because the information for linking a chromosome is missing. Only if at least one non-tetraplotig coverage marker can be grouped, the tetraploitig marker of the contigs can be grouped.

Meiotic recombination can influence the linkage grouping. In the most simple case, i.e., when no pollen genome carries a single meiotic recombination event, all PAPs at coverage markers from the same haplotype would be identical while PAPs at coverage markers from other haplotypes would be different (because haplotypes randomly occur in pollen genomes in a pair-wise manner), and thus they can be easily grouped into haplotypes. In the presence of meiotic recombination, a few crossovers along each chromosome change the PAP values of the coverage markers, but as recombination is rare the PAP values change only marginally. Thus, linkage grouping based on PAPs works even in the presence of meiotic recombination.

#### Haplotype-specific PacBio HiFi read separation and haplotype assembly

HiFi reads were classified into 48 groups based on alignments to the 50 kb coverage markers. Specifically, to assign a read to a marker, at least 500 bp of the read had to be aligned to the marker (reads aligning two neighboring markers were assigned to the marker with a larger overlapping size). Reads overlapping non-haplotig marker were randomly assigned to one of the the marker-associated groups.

Each set of HiFi reads was independently assembled using *hifiasm* (version 0.7) with default settings. The resulting contigs were first polished with short reads using *pilon* with *--fix bases --changes --diploid --mindepth* 0.8 and then with HiFi reads using *racon*[12] with *-u --no-trimming*.

#### Analysis of HiFi reads not assigned to any of the 48 haplotypes

The non-grouped reads were provided to *hifiasm* to perform independent assembly under default settings. All raw contigs were aligned to the NCBI nucleotide database using *blastn*[13] under default settings. The top blasting hit was selected to define whether a contig was from organellar genomes, and the percentage overlapping organelle genomes was calculated as the aligned length divided by the total contig length (Figure S7).

#### Evaluation of haplotyping accuracy

For each haplotype assembly and the sequencing data of the parental genomes, *k*-mers (*k*=21) were counted using *KMC*[14]. Specifically, *k*-mers found in *Hera* but not in *Stieglitz* (with a coverage of 6-12), as well as *k*-mers found in *Stieglitz* but not in *Hera* (with a coverage of 5-11) were selected using *kmc_tools simple*. For each haplotype, the set of assembled *k*-mers were intersected with the two sets of parental-specific *k*-mers (using *kmc_tools simple* with sub-function *intersect*), which revealed *k*-mers common with either of the parental genomes. As a haplotype can only be inherited from one of the parents, it is expected to find parental-specific *k*-mers only of one parent. The overall haplotyping precision was determined as the total number of correctly phased *k*-mers divided by the total number of *k*-mers investigated in the 48 haplotype assemblies. Note, this was done before and after contig polishing, where we observed the same haplotyping accuracy.

#### Haplotype-specific Hi-C read separation

To avoid mis-joined haplotypes (which is a common problem in haplotyping and scaffolding with chromosome conformation capture data), we removed Hi-C read pairs linking different haplotypes. All Hi-C reads were first aligned to the initial assembly (of 6,366 contigs) using *bowtie2*. If a read pair can be aligned to coverage markers from a single group, then it was assigned to that group, otherwise the read pair was removed from analysis. Note that when a read pair could be aligned to coverage markers of multiple groups simutaneouly, it was randomly assigned to one of the groups. This process led to 70x per haplotype out of the raw coverage of 130x per haplotype.

#### Haplotype-specific contig scaffolding using group-specific Hi-C reads

Each haplotype-specific contig-level assembly was indexed with *bwa index* (with *-a bwtsw*) (version 0.7.15-r1140)[15] and *samtools faidx*. The haplotype-specific Hi-C read pairs were aligned using *bwa aln* and *bwa sampe*. Aligned reads (in pairs) were converted into BAM files using *samtools view* with options of *-b -F12*. The BAM files were filtered with *filterBAM_forHiC.pl* (from *ALLHiC* package, version 0.9.13)[16] to remove non-uniquely mapped reads. Then BAM files were converted to bed files using *bamToBed* (from *bedtools* package) and sorted by read name. The bed files were provided to *SALSA2* (run with -*s* 100000000 -*m yes* -i *10* -e *DNASE*)[17]. Potential chimeric contigs were broken at the chimeric sites given by *SALSA2* output file of *input_breaks*, leading to a new set of contigs for each of the 48 original groups.

For each new group of contigs, the above process of contig indexing, Hi-C read alignment and BAM filtering was repeated. Then, for each haplotype, *ALLHiC_partition* was run with *-e GATC -k 1 -m 25*; *allhic extract* was run with --*RE GATC*; *allhic optimize* and *ALLHiC_build* were run with default settings; the chromosome contact map was visualized with *ALLHiC_plot* at 1 Mb resolution, where obvious mis-placement/orientation of large contigs were visually identified and manually corrected (Figure S8).

#### Evaluation of assembly quality

The trimmed short reads from the two 10x single cell CNV libraries (DNA) were used to create a *k*-mer databse (*otava_genome.meryl*) for the 'Otava' genome using *meryl*[18] with options "*k=21 count threads=4 memory=8g*". Then the fully combined set of *Hera* 1,2 and *Stieglitz* 1,2 sequences (*otava_genome.fa*) were compared with the *k*-mer database (*otava_genome.meryl*) to investigate the completeness (97.3%) and the base accuracy (QV>51.7) of the final assembly, using "*merqury.sh genome.meryl otava_genome.fa full_genome*"[18].

#### Comparison of chromosome sequences

Within each of the twelve homologous LGs, the chromosome-level sequences of the four haplotypes were aligned to each other as well as to the recently assembled *DM* genome using *minimap*2 with *-ax asm20 --eqx*. For each pair of haplotypes, the alignments were provided to *SyRI*[19], which searched for synteny, single-nucleotide level differences as well as large-scale structural variations (with *-k -F S*).

#### Identification of identical-by-decent blocks

The initial 50 kb coverage markers were aligned to the chromosome-level haplotype-resolved assembly using *minimap2*, where a maximum of 2,3,4 hits were allowed for diplotig, triplotig and tetraplotig related markers. Neighbouring (aligned) markers of the same type were connected into larger regions if their distance was less than 200 kb.

#### Identification of potentially collapsed variants within IBD blocks

All short reads of the two 10x single-cell CNV libraries were aligned to the polished full assembly using *bowtie2*. Bed files were created with coordinates of IBD blocks shared by two, three, and four haplotypes, and short reads aligned to these regions were extracted using *bedtools intersect.* For all groups of IBD blocks, a single-copy of the shared sequence from each group was collected, which was subsequently combined into one fasta file to create a set of sequences as reference, which was indexed by *bowtie2*. The extracted short reads were aligned to the created reference sequences using *bowtie2*. Variants were called using *bcftools*. The results were collected in Data S7, where it showed a maximum of 1 SNP per 72 kb could have been missed from phasing in IBD blocks.

#### Genome annotation and assessment

Protein-coding genes for each haplotype chromosome were annotated with three types of evidences, including *ab initio* gene predictions (considering outputs by *Augustus*[20], *GlimmerHMM*[21] and *SNAP*[22]), transcripts assembled from Illumina short RNA-seq reads, and alignments of homologous protein sequences. Specifically, protein sequences from *Solanum tuberosum L.*[10,23], *Arabidopsis thaliana* and other plant proteins from UnitProtKB were aligned to each haplotype assembly independently using *Exonerate*[24] with options of *--percent* 60 *--minintron* 10 *--maxintron* 60000. RNA-seq reads from two recent potato genome assembly work[10,23] were downloaded. All reads were first aligned to the full haplotype-resolved genome assembly using *HISAT2* version 2.2.0[25]. Then for each of the 12 linkage groups, reads aligned to the respective four haplotypes were extracted and combined as one set with *samtools view -L* and *bedtools bamtofastq*. Within each linkage group, each set of reads from the group were re-aligned to each haplotype sequence independently using *HISAT2*, and transcripts were assembled using *StringTie*[26]. Finally, all the above evidences were integrated with *EvidenceModeler*[27] in order to generate high-quality gene models for each haplotype assembly. Transposon elements (TE) were annotated using *RepeatModeler* and *RepeatMasker* (http://www.repeatmasker.org). TE related genes were filtered by investigating their overlapping with TEs (overlapping percent > 30%), sequence alignment with TE-related protein sequences and *A. thaliana* TE related gene sequences (requiring *blastn* alignment identity and coverage larger than 30%).

To rescue potentially mis-annotated genes, all gene models were further improved. Specifically, against each of the four haplotypes, we first aligned the gene sequences of the other three haplotypes using *blastn*, and similarly, aligned the protein sequences of the other three haplotypes against those from the target haplotype using *blastp*[13] and *Scipio*[28]. If there were counterparts in the target haplotype to the other three (based on the alignments), the potential missing genes were added according to gene models given by *ab initio* prediction. Besides, gene models were split into smaller genes or merged as larger genes if all the alignments of genes from the other three haplotypes indicated a mis-merged or mis-split gene model.

The final assembly and annotation completeness were evaluated by *BUSCO* (version 4.1.4)[29] with 2,326 single-copy genes from the lineage database "eudicots_odb10". The functional annotation of genes was performed with *InterProScan* (version 5.48)[30] with default parameters except for option "-goterms". The GO terms were extracted for GO functional enrichment analysis by the R package *ClusterProfiler*[31]. The noncoding RNA was annotated with the tool *Infernal* v1.1[32] by searching the database Rfam v14.3[33]. The ajacent rDNAs with distance smaller than 5 kb were clustered as the potential rDNA clusters.

#### Characterization of noncoding RNA

First, all transcripts were assembled using *StringTie* (v1.3.4d) with default parameter settings using our RNA-seq data and public RNA-seq data. Transcripts overlapping with the exons of annotated protein-coding gene at the sense strand or shorter than 200 bp were removed with the script *FEELnc_filter.pl* in the tool *FEELnc*[34]. Second, the transcripts with a potential for protein-coding were filtered by the tool *CPC2*[35]. Third, we used the *blastx* to align the filtered transcripts against the protein sequences from Swiss-Prot database[36] to filter strong hits with some cutoffs (alignment identity > 35, alignment length >40 aa and alignment coverage of the query or subject sequence >35) following a previous study[37].

#### Gene family and PAV analysis

We clustered the protein sequences from all four haplotypes using *OrthoFinder* (version 2.2.6)[38] with default parameters. The gene presence/absence variants between the haplotypes were identified based on the gene groups resulting from *OrthoFinder*. To decrease the false absence of genes due to some residual missing genes in annotation, the focal haplotype was predicted to also contain a homolog of a gene annotated in other haplotypes when the gene was aligned to the focal haplotype with a high alignment coverage (>90%) and identity (>90%)

and no loss-of-function variations (based on the variations calling from the tool *SyRI* as described above).

#### Crossover detection (at LG 4) in pollen genomes

For each of the four haplotype sequences at LG 4, 30x error-free Illumina paired-end reads were simulated using *pirs*[39] with options of *-m* 300 *-v* 10 *-l* 100 *-x* 25 *-e* 0 *-a* 0 *-g* 0. The reads were aligned to *Chr-4.Stie_2* as reference sequence using *bowtie2* under default settings, and variations were called using *bcftools* (version 1.9). For each of the non-reference sequences, we selected variations specific to all other haplotypes and with a minimum mapping quality of 150, minimum allele frequency of 0.99 and a coverage of the alternative allele in [16, 34]. For the reference chromosome, if all the other three haplotypes had the same allele but different from the reference, the allele would be selected as specific in the reference chromosome. Such variations were filterted for minimum mapping quality of 150, minimum allele frequency of 0.99 and a coverage of the alternative allele in [16, 34]. All these haplotype-specific variations were used as SNP markers in genotyping pollen genome sequencings.

Each read set of the 717 pollen nuclei sequencing was respectively aligned to the *Chr-4.Stie_2* sequence, and consensus were called using *bcftools*. The allele read counts were obtained at the above-defined SNP markers for chromosomes 4, based on which crossovers were detected.

#### Allelic expression analysis

*Quality control*. Short reads from RNA sequencing were trimmed with *Trimmomatic*[40] (version 0.39) under paired-end mode, with options "*ILLUMINA:adapters.fa:2:30:10:8:true (adapters provided by the tool itself) SLIDINGWINDOW:4:15 LEADING:3 TRAILING:3 MINLEN:36*".

*Read separation*. All reads were aligned to the final haplotype-resolved assembly using *HISAT2* version 2.2.0 with option "*-k 1*", and reads were thus separated into 48 haplotype-specific groups with *samtools view -L* and *bedtools bamtofastq*.

*Expression analysis using Stieglitz 1 genome as reference.* Within each of the 12 linkage groups, the four sets of haplotype-wise RNA-seq reads were independently aligned to the respective *Stieglitz* 1 chromosome as reference. The number of fragments at each gene from each chromosome was quantified using *HTSeq*[41] version 0.13.5 with options "*--mode=union --nonunique=none --secondary-alignments=ignore --stranded=no*" (given the gff file for that chromosome). For testing dominance in allelic expression and effect of allelic copy number on expression, the fragment counts were normalized as fragments per kilobase per million reads (FPKM) and log2-scaled. The genes with four allelic copies were selected in analyzing differential allele expression, following the existing pipeline[42], where the log2-

transformed CPM (count per million reads) at each gene was used for measuring the allele expressen level. After excluding genes with less than 3 replicates showing CPM over 1.0, 11,154 expressed genes were kept. Paired comparions on expression of the four alleles were performed, and 1,219 were found to be differentially expressed in at least one pair of haplotypes with adjusted *p*-value<0.05 (Data S11).

Note that the well clustering of the three technical replicates regarding four haplotypes based on the haplotype-specific expression of all genes from *Stieglitz-1* genome (using *hclust* in R) showed that there was a high consistency between the replicates (Figure S28), thus the final FPKM value at each gene was taken as the average of the values of the three replicates in all related analysis (except for the procedure of differential analysis where replicates were not merged). At a gene, for four alleles with FPKM values of *x={x1,x2,x3, x4}*, the *z*-score (in heatmaps) for each allele *i* (in {1,2,3,4}) was calculated as *(xi - mean(x)) / sd(x)*, with *mean(x)* giving the average of *x* and *sd(x)* giving the standard deviation of *x*.

#### Methylation analysis

*Quality control*. Short reads from methylation sequencing were trimmed with *Trimmomatic* (version 0.39) under paired-end mode, with options "*ILLUMINA:adapters.fa: 2:30:10:8:true (adapters provided by the tool itself) SLIDINGWINDOW:4:15 LEADING:3 TRAILING:3 MINLEN:36*".

*Read separation*. All reads were first aligned to the initial assembly (of 6,366 contigs) using *bismark*[43] version 0.23.0 (with options "*--hisat2 --score_min L,0,-0.6*"), to avoid the default masking of reads from IBD regions. If a read pair can be aligned to coverage markers linking a single haplotype group, then it was assigned to that group. If a read pair could be aligned to coverage markers linking multiple groups, it was randomly assigned to one of the groups.

*Methylation calling at 48 haplotype-specific chromosomes*. The reads assigned to each haplotype-specific group were independently re-aligned to the respective haplotype-resolved chromosome using *bismark* under the same options plus "*--non_directional*" (which gave alignments to all four bisulfite strands). Each bam file was deduplicated using *deduplicate_bismark*, after which methylation at CG, CHG and CHH contexts were simultaneously called with *bismark_methylation_extractor* with options "*--comprehensive --cytosine_report --CX --bedGraph*". Given a specific genomic region/window, the methylation level at CG(/CHG/CHH) context was determined as *C_count/(C_count+T_count)*, where *C_count* was the number of reads carrying the methylated cytosine in CG(/CHG/CHH) context and *T_count* was the number of reads carrying un-methylated cytosine in CG(/CHG/CHH) context. Following this, a sliding window based quantification of the methylation level were performed along the 48 chromosomes (window size: 2 Mb, step: 50 kb; main text Figure 2b).

*Methylation calling at 12 Stieglitz-1 chromosomes as references*. Within each of the linkage groups, the four haplotype-wise reads were independently aligned to the respective *Stieglitz*-1 chromosome as reference using *bismark* under the same options (as above) plus "--*non_directional*" (which gave alignments to all four bisulfite strands). The methylation calling was done in a similar way as given above. The results were used to perform correlation analysis with allele-specific expression (main text Figure 4g).

*Comparison of the level of methylation between IBD blocks and the corresponding regions in synteny in homologous haplotypes.* IBD blocks (at 50 kb resolution) that were shared by two or three haplotypes were investigated. According to the synteny between haplotypes (from *SyRI* based analysis), the counterparts of such IBDs were located in the haplotypes being homologous to the haplotypes showing the IBDs. The counterparts were re-assigned to IBD groups if they were harboured by any known IBD blocks. This led to two sets, one with IBD blocks and the other with regions (from other haplotypes but) in synteny with the IBD blocks. Methylation level at CG(/CHG/CHH) context was calculated for each block within the two sets, and *t*-test was used to investigate the difference between the two sets (Figure S27).

Note that, using the 12 *Stieglitz-1* chromosomes as reference, the well clustering of the three technical replicates regarding four haplotypes based on either CG, CHG or CHH methylation level in 50 kb windows at a step of 25 kb (using *hclust* in R) showed that there was a high consistency between the replicates (Figure S28), thus the final methylation level (and the related *z*-score, similarly calculated as given in gene expression analysis) at each window was taken as the average of the levels at three replicates in all related analysis.

#### Supplementary Figures

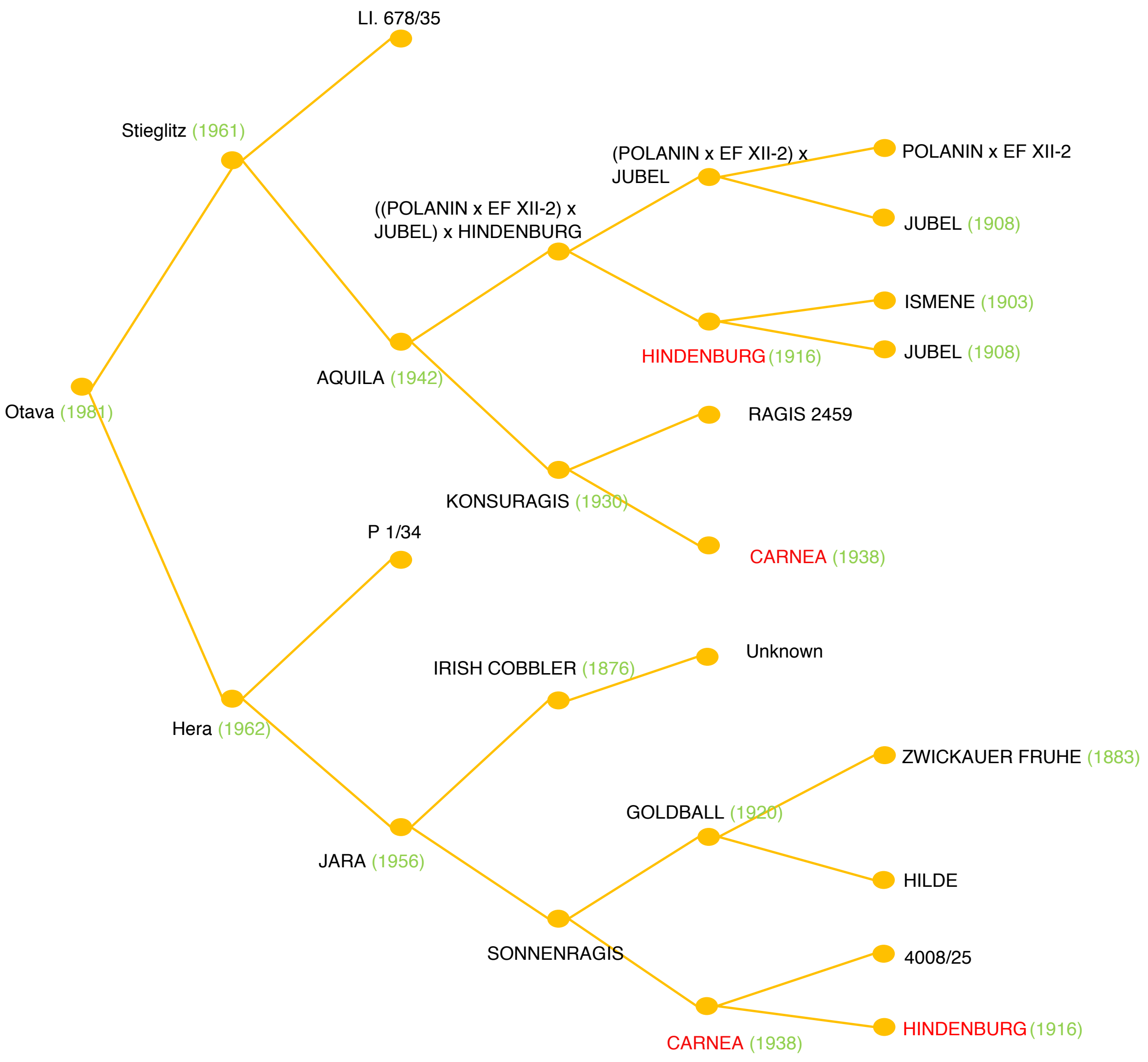


**Figure S1. Pedigree of 'Otava'.** The potato cultivar 'Otava' resulted from a cross of 'Hera' and 'Stieglitz' in 1981. Note that 'Hera' and 'Stieglitz' have common ancestors like 'HINDENBURG' (1916) and 'CARNEA' (1938) explaining the presence of IBD regions between the four haplotypes of *Otava* genome. This illustration was modified from the *Potato Pedigree Database (https://www.plantbreeding.wur.nl/PotatoPedigree)*[44,45].

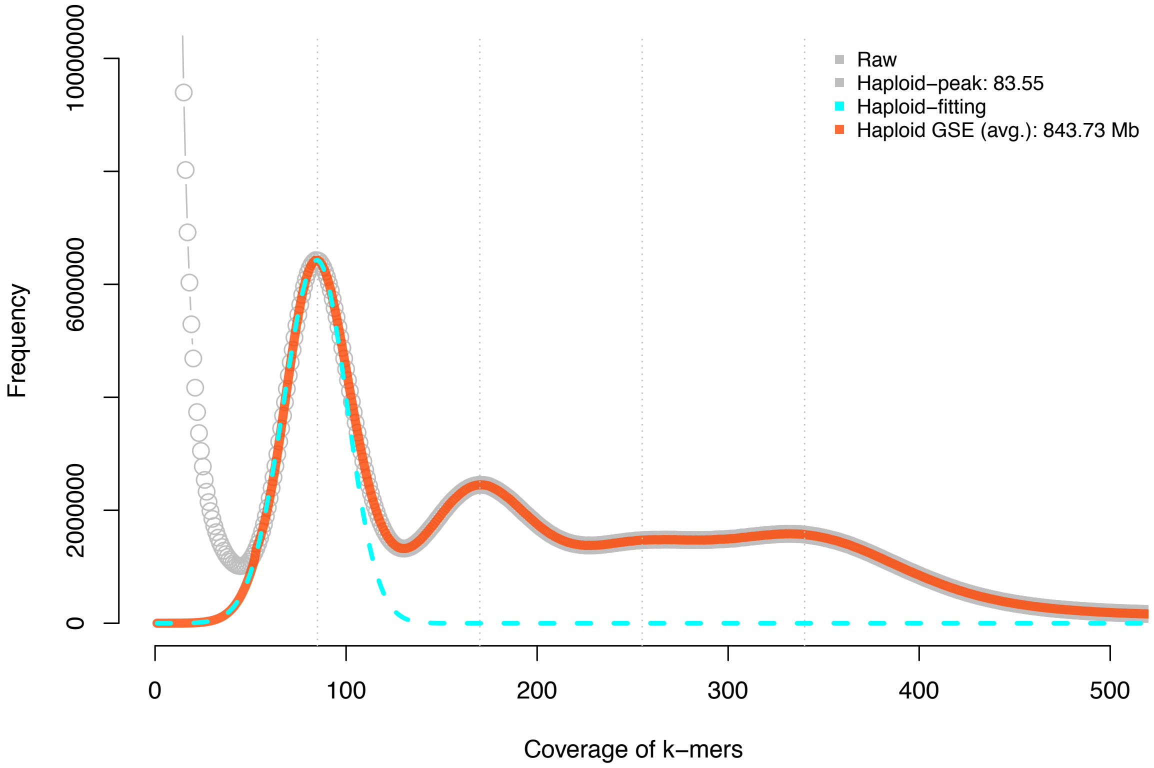


**Figure S2.** ***K*-mer distribution of the tetraploid genome of 'Otava' and *k*-mer based genome size estimation.** Note, 844 Mb depicts the estimated haploid genome size, while the tetraploid genome size would be four times the haploid genome size, i.e., 3,375 Mb.

a

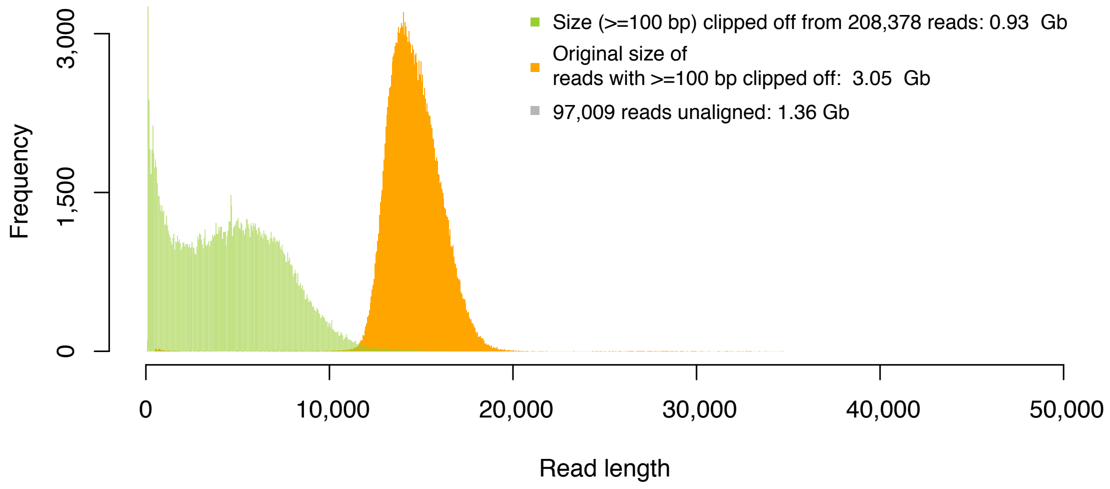


b

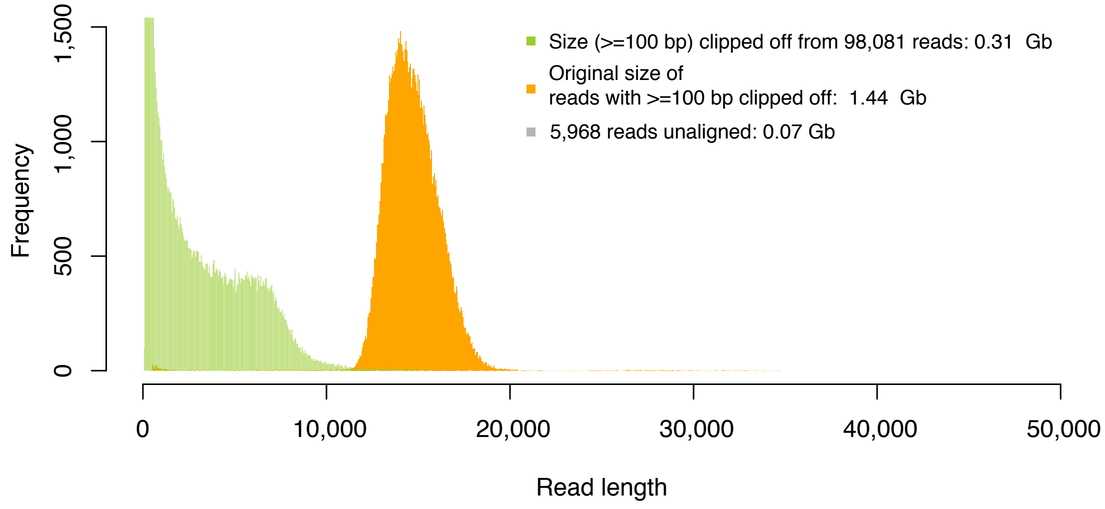


**Figure S3. Aligning (102.2 Gb) PacBio HiFi reads to the raw and the initial assemblies (after purging low-coverage contigs). a.** Aligning HiFi reads to the raw assembly of 17,153 contigs led to 208,378 reads, which were clipped with >=100 bp (0.93 Gb bases clipped off) and 97,009 reads (1.36 Gb), which were not aligned. This gave an overall alignment rate of 97.8% at base level. **b.** Aligning HiFi reads to the (purged) initial assembly of 6,366 contigs (with an N50 of 2.1 Mb) led to 98,081 reads, which were clipped with >=100 bp (0.31 Gb bases clipped off) and 5,968 reads (0.07 Gb), which were not aligned. This gave an overall alignment rate of 99.6% at base level. The increased alignment rate supported that the purging had removed redundant information, for example, repeated representation of some genomic regions which could be induced by sequencing errors in HiFi reads, from the raw assembly. (*x*-axis in bp)

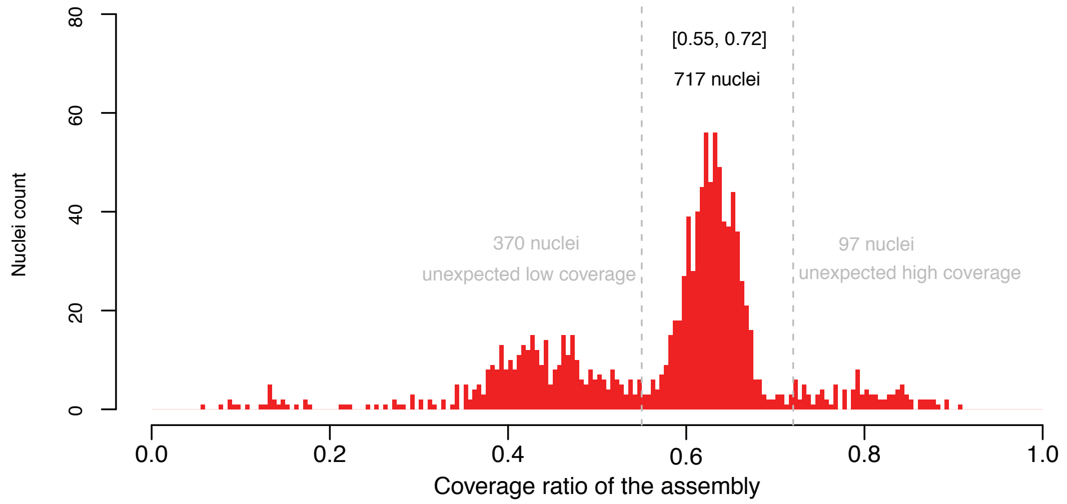


**Figure S4. Histogram of the coverage ratio of each pollen after aligning the respective reads against the initial assembly (of 6,366 contigs).** Initially, 1,184 pollen nuclei (with a minimum of 40,000 151 bp read pairs) were extracted from the two 10x single cell CNV libraries. By aligning each read set to the initial assembly, we found a major cluster containing 717 nuclei that could cover the assembly with a ratio of 0.55 to 0.72 and selected them to perform linkage grouping, where the minimum read pair number was 70,356, overall leading to a mean coverage of 0.18x of the haploid genome.

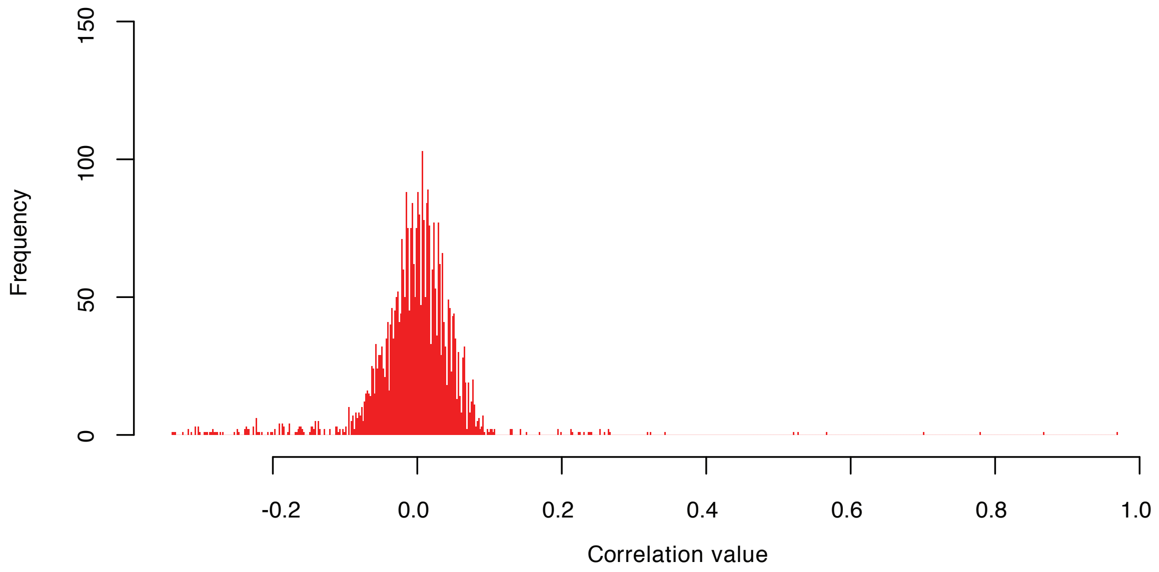


**Figure S5. Distribution of the PAP-defined correlation values of an arbitrarily selected representative coverage marker of one contig with the coverage markers of all other contigs.** Based on this distribution, a minimum correlation value of 0.55 was selected as threshold to identify marker pairs, which were physically closely located. A maximum correlation value of -0.25 was selected to support that two markers could be from (neighboring) allelic regions on homologous chromosomes.

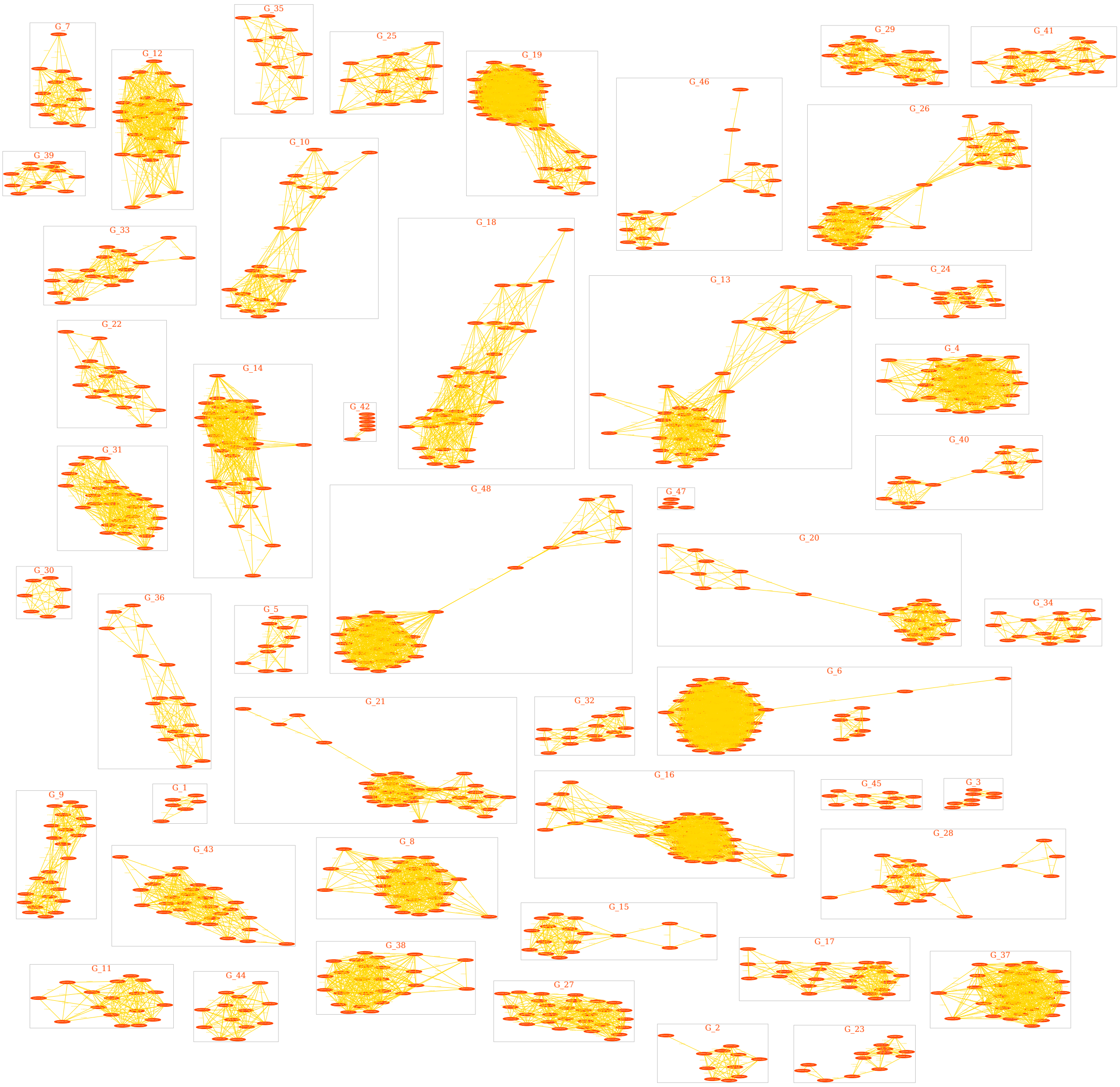

**Figure S6. The graph (consisting of 48 sub-graphs) visualizes the clustering of haplotigs of over 100 kb into 48 groups representing the 48 haplotype-specific chromosomes.** Vertices in red represent contigs, edges in yellow represent a positive correclation (over 0.55) of the PAPs of the connected coverage markers (vertices). All nodes in the same box in gray represent a group of contigs belonging to the same chromosome labeled with G_*i*, where *i*=1, 2, …, 48. The graph was visualized with *Graphviz* (https://graphviz.org/).

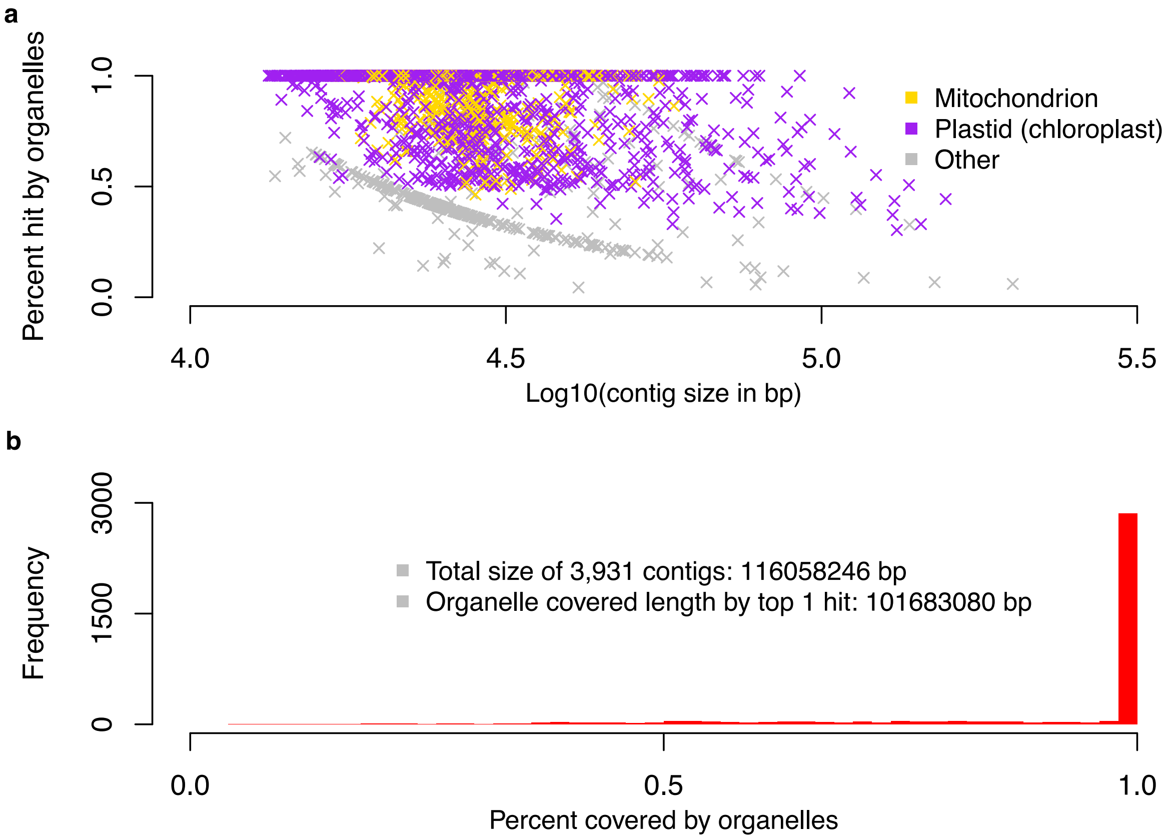


**Figure S7. Alignment of contigs (that were assembled from 9.9 Gb non-grouped HiFi reads) to the NCBI nucleotide database.** Among all 3,931 contigs (with a total size of 116 Mb), 3,439 could be at least partially aligned to organelle sequences (**a**), among which 2,814 contigs fully overlapped organelle sequences (**b**). At base-level, 101.7 Mb sequences could be aligned to organelle genomes, accounting for nearly 88% of the total contig size of 116 Mb.

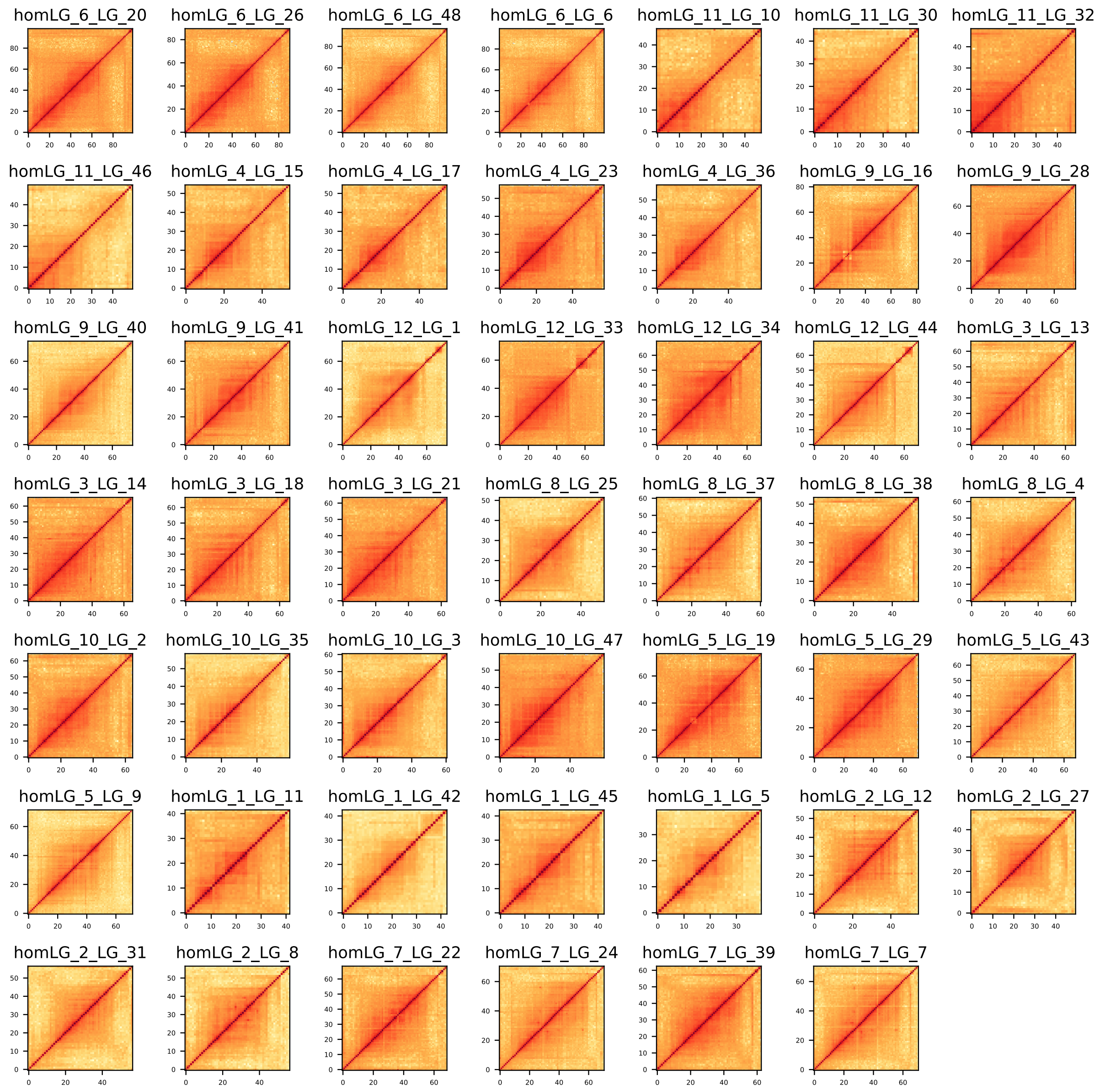


**Figure S8. Hi-C contact map for each of the 48 haplotype-specific chromsomes.** *HomLG* 6, 11, 4, 9, 12, 3, 8, 10, 5, 1, 2, 7 correspond to LG 1 to 12 of *DM*[9,10]. For example, *homLG_6* corresponds to LG 1 in the *DM* assembly. The four sub-groups (*LG_20*, *LG_26*, *LG_48* and *LG_6*) correspond to the four haplotype-specific chromosomes of 'Otava' (for which the identifiers were defined by the linkage grouping step of gamete binning as given by Figure S6).

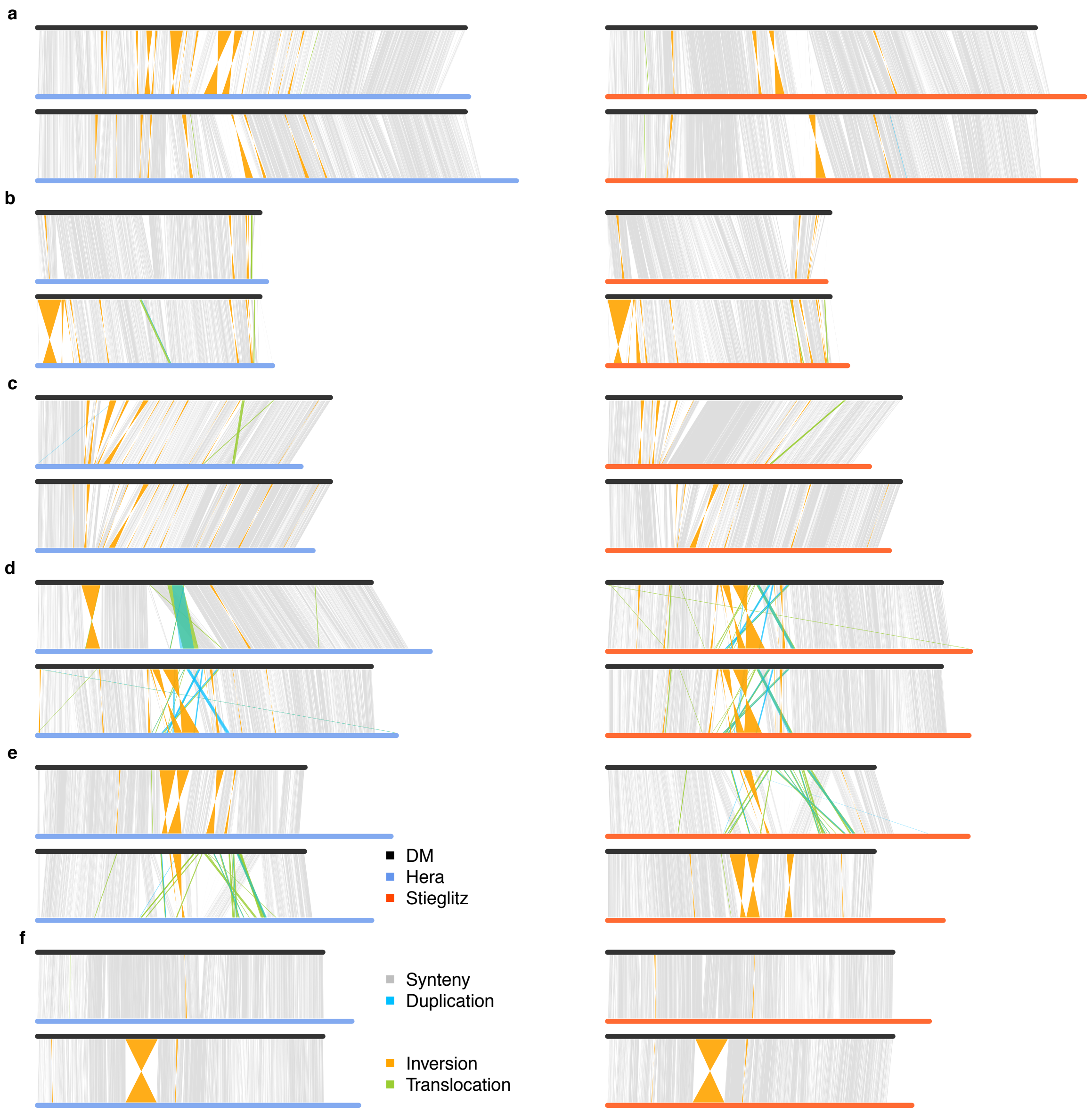


**Figure S9. Comparison of LG 1-6 of the 'Otava' assembly to the *DM* assembly[9,10]. a-f**: LG 1-6. Within each LG, there were four haplotype sequences from 'Otava' (with two of them inherited from 'Hera' (in light blue) and two inhertited from 'Stieglitz' (in light red) ). Alignments between any two sequences above 50 kb are shown.

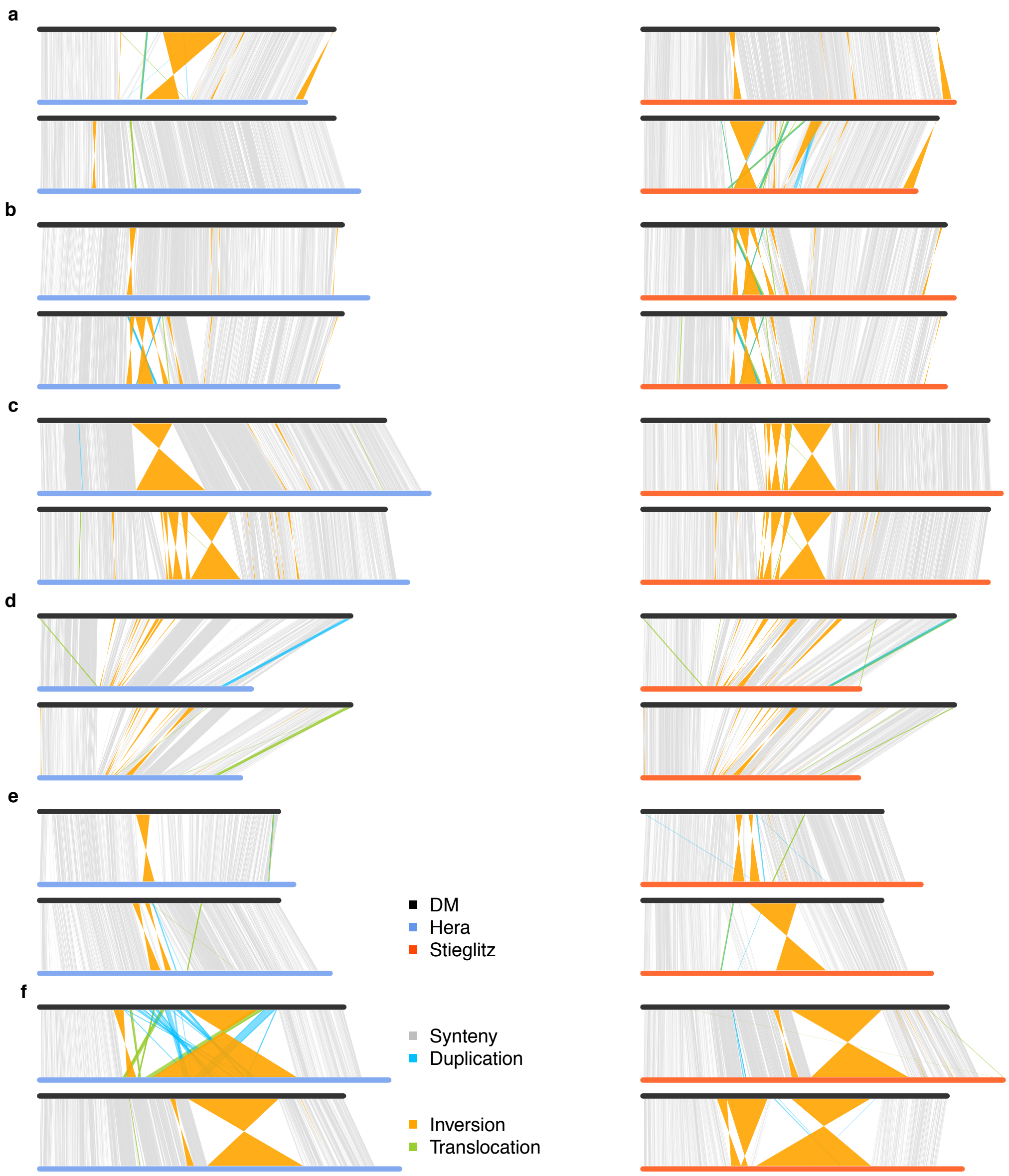


**Figure S10. Comparison of LG 7-12 of the 'Otava' assembly to the *DM* assembly[9,10]. a-f**: LG 7-12. Within each LG, there were four haplotype sequences from 'Otava' (with two of them inherited from 'Hera' (in light blue) and two inhertited from 'Stieglitz' (in light red)). Alignments between any two sequences above 50 kb are shown. Speficifically, for LG-10 (given by d), the four haplotype-specific chromosomes showed five large gaps when aligned to the DM chromosome, incating that there were potential chromosomal large rearrangements between the two cultivars.

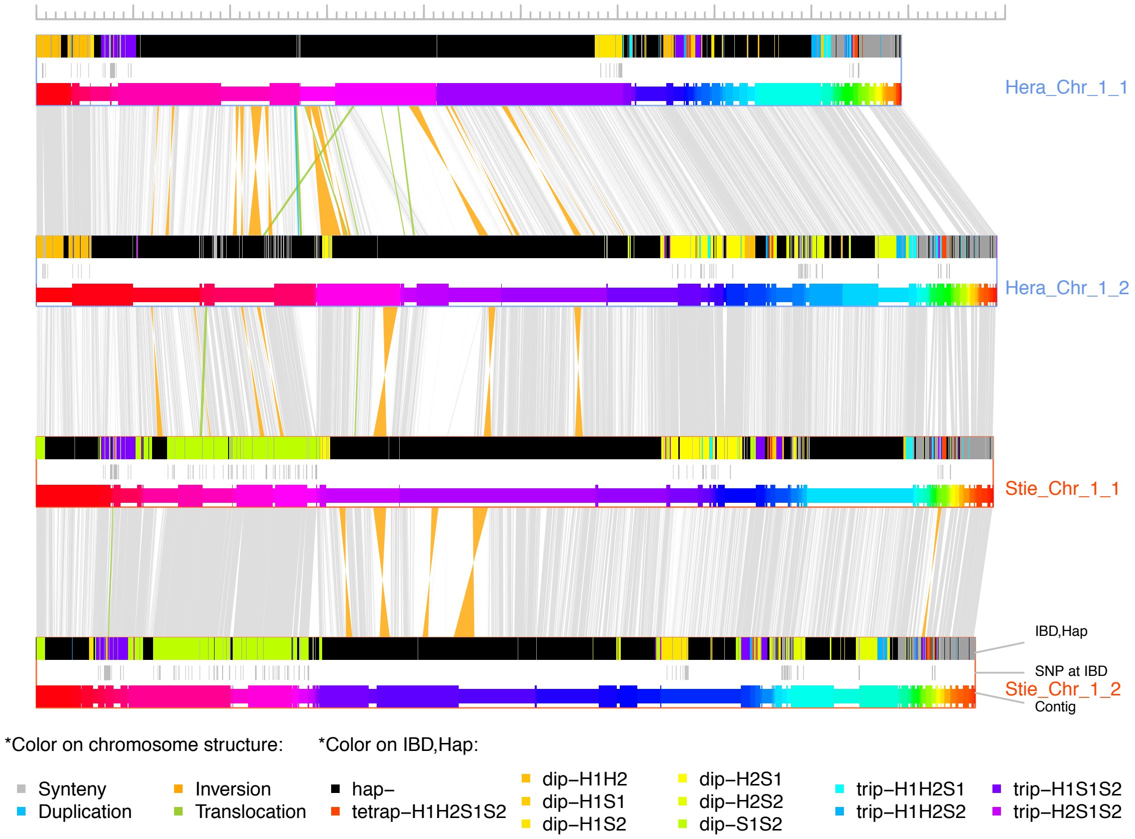


**Figure S11. Comparison of LG-wise haplotypes at LG 1**. Each of the four horizontal rows represents a haplotype of the chromosome (*Hera* 1, *Hera* 2, *Stieglitz* 1 and *Stieglitz* 2). Three types of information are within each haplotype box, top: IBD and unique regions along the chromosome (at 50 kb resolution), middle: distribution of SNPs at IBD regions, and bottom: scaffolded contigs. For each chromosome, three pair-wise structural comparisons are shown highlighting inversions (orange), duplications (blue), translocations (green) and syntenic regions (grey). Note that all the regions involving the breakpoints of the SVs ended within contigs. IBD regions can be shared by two (dip), three (trip) or even four (tetrap) haplotypes. The x-axis scale: 0-100 Mb.

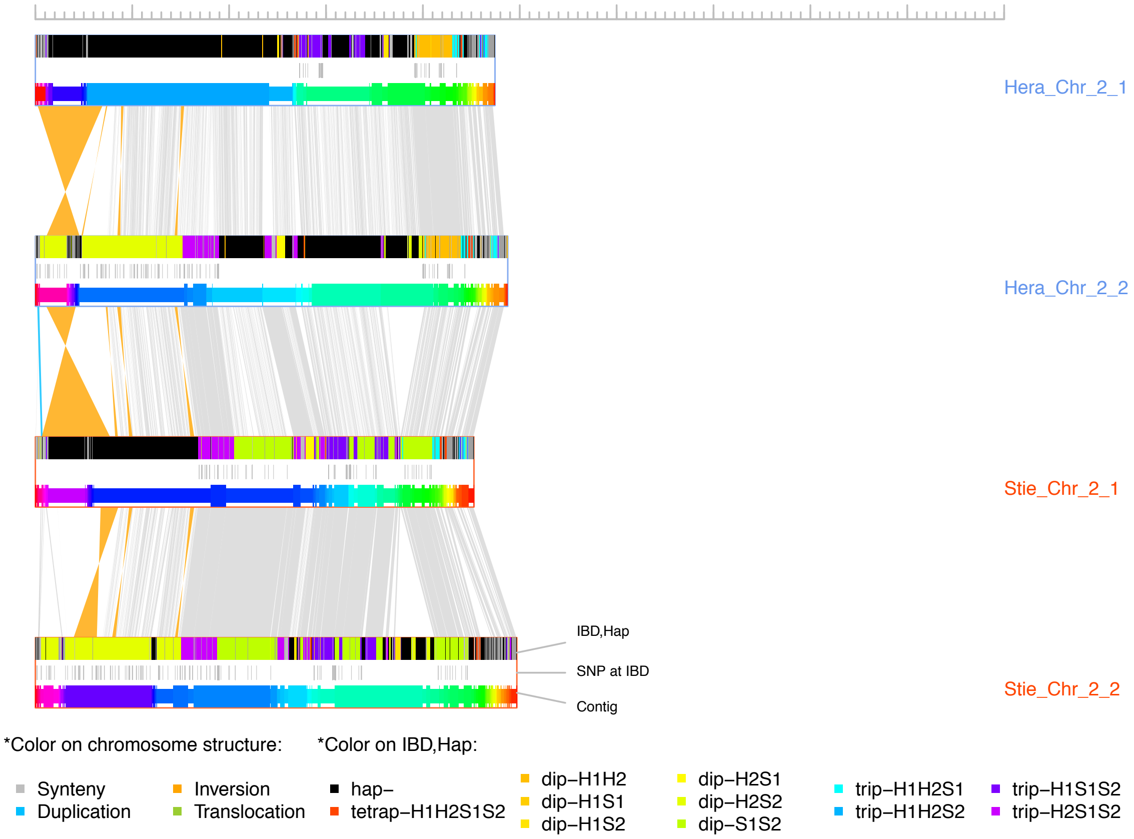


**Figure S12. Comparison of LG-wise haplotypes at LG 2**. Each of the four horizontal rows represents a haplotype of the chromosome (*Hera* 1, *Hera* 2, *Stieglitz* 1 and *Stieglitz* 2). Three types of information are within each haplotype box, top: IBD and unique regions along the chromosome (at 50 kb resolution), middle: distribution of SNPs at IBD regions, and bottom: scaffolded contigs. For each chromosome, three pair-wise structural comparisons are shown highlighting inversions (orange), duplications (blue), translocations (green) and syntenic regions (grey). Note that all the regions involving the breakpoints of the SVs ended within contigs. IBD regions can be shared by two (dip), three (trip) or even four (tetrap) haplotypes. The x-axis scale: 0-100 Mb.

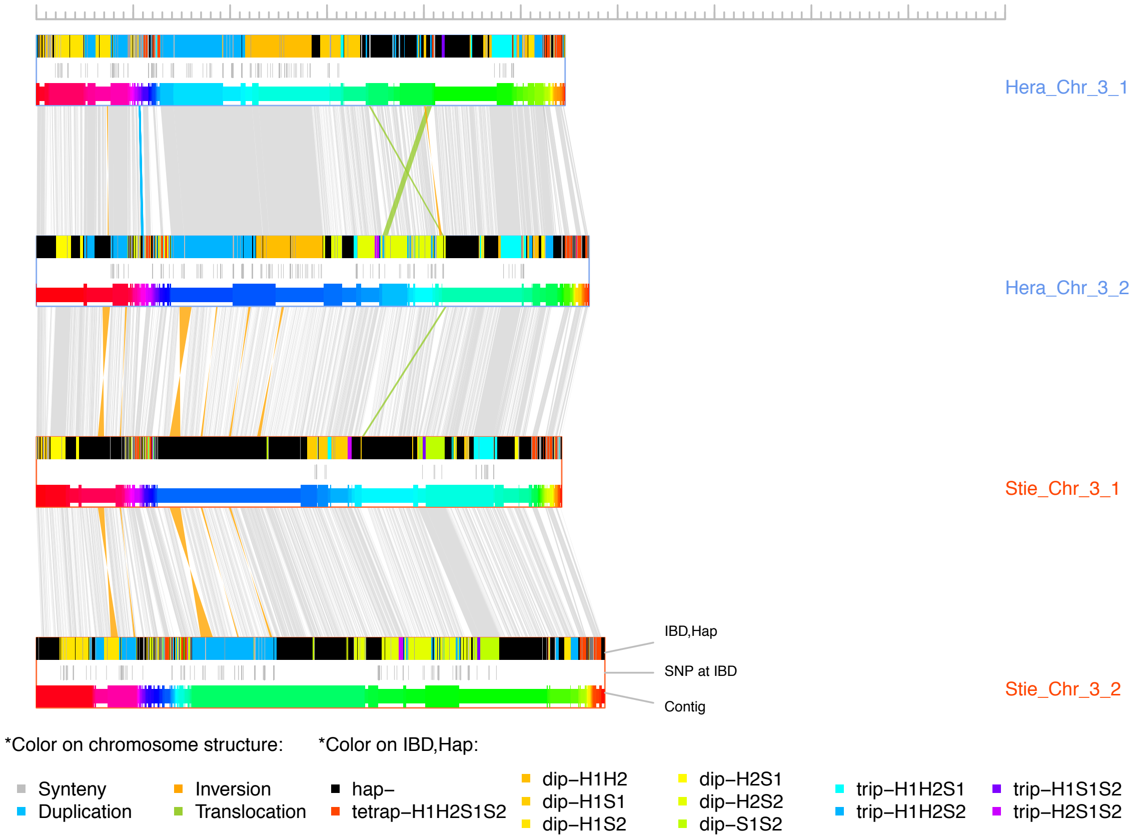


**Figure S13. Comparison of LG-wise haplotypes at LG 3**. Each of the four horizontal rows represents a haplotype of the chromosome (*Hera* 1, *Hera* 2, *Stieglitz* 1 and *Stieglitz* 2). Three types of information are within each haplotype box, top: IBD and unique regions along the chromosome (at 50 kb resolution), middle: distribution of SNPs at IBD regions, and bottom: scaffolded contigs. For each chromosome, three pair-wise structural comparisons are shown highlighting inversions (orange), duplications (blue), translocations (green) and syntenic regions (grey). Note that all the regions involving the breakpoints of the SVs ended within contigs. IBD regions can be shared by two (dip), three (trip) or even four (tetrap) haplotypes. The x-axis scale: 0-100 Mb.

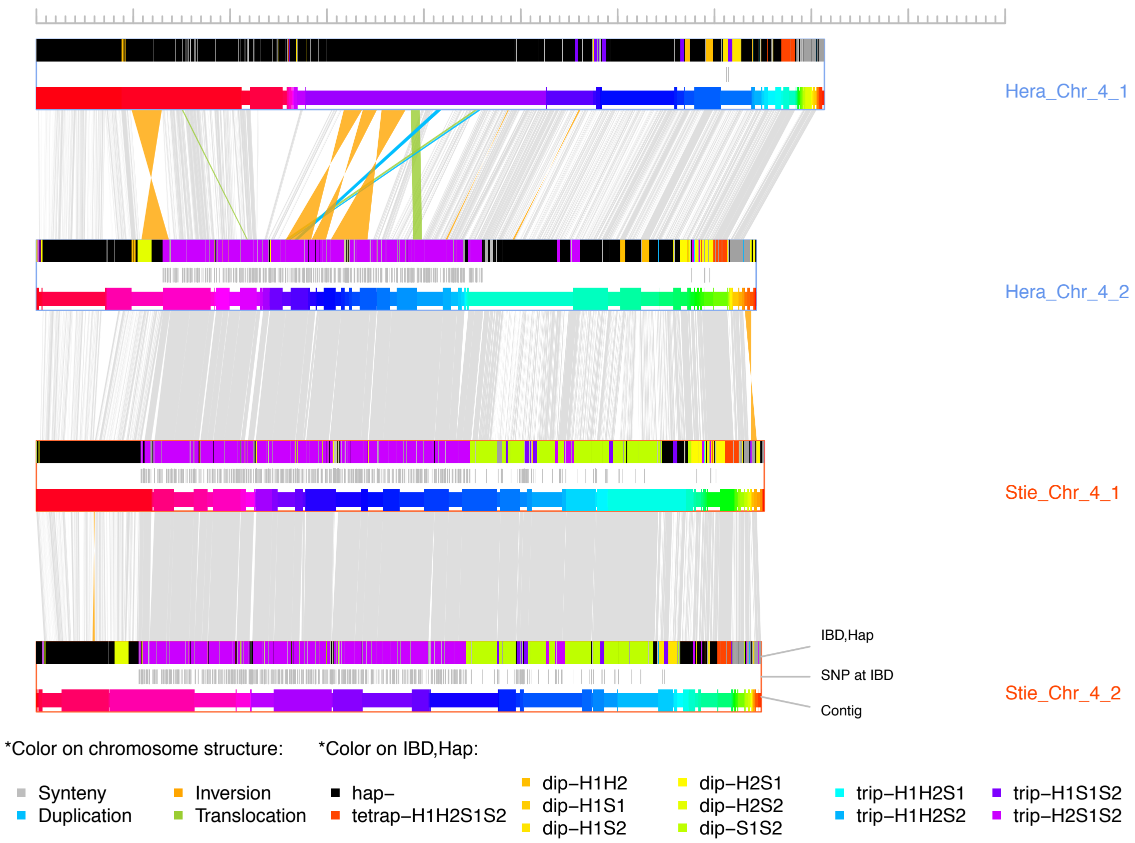


**Figure S14. Comparison of LG-wise haplotypes at LG 4**. Each of the four horizontal rows represents a haplotype of the chromosome (*Hera* 1, *Hera* 2, *Stieglitz* 1 and *Stieglitz* 2). Three types of information are within each haplotype box, top: IBD and unique regions along the chromosome (at 50 kb resolution), middle: distribution of SNPs at IBD regions, and bottom: scaffolded contigs. For each chromosome, three pair-wise structural comparisons are shown highlighting inversions (orange), duplications (blue), translocations (green) and syntenic regions (grey). Note that all the regions involving the breakpoints of the SVs ended within contigs. IBD regions can be shared by two (dip), three (trip) or even four (tetrap) haplotypes. The x-axis scale: 0-100 Mb.

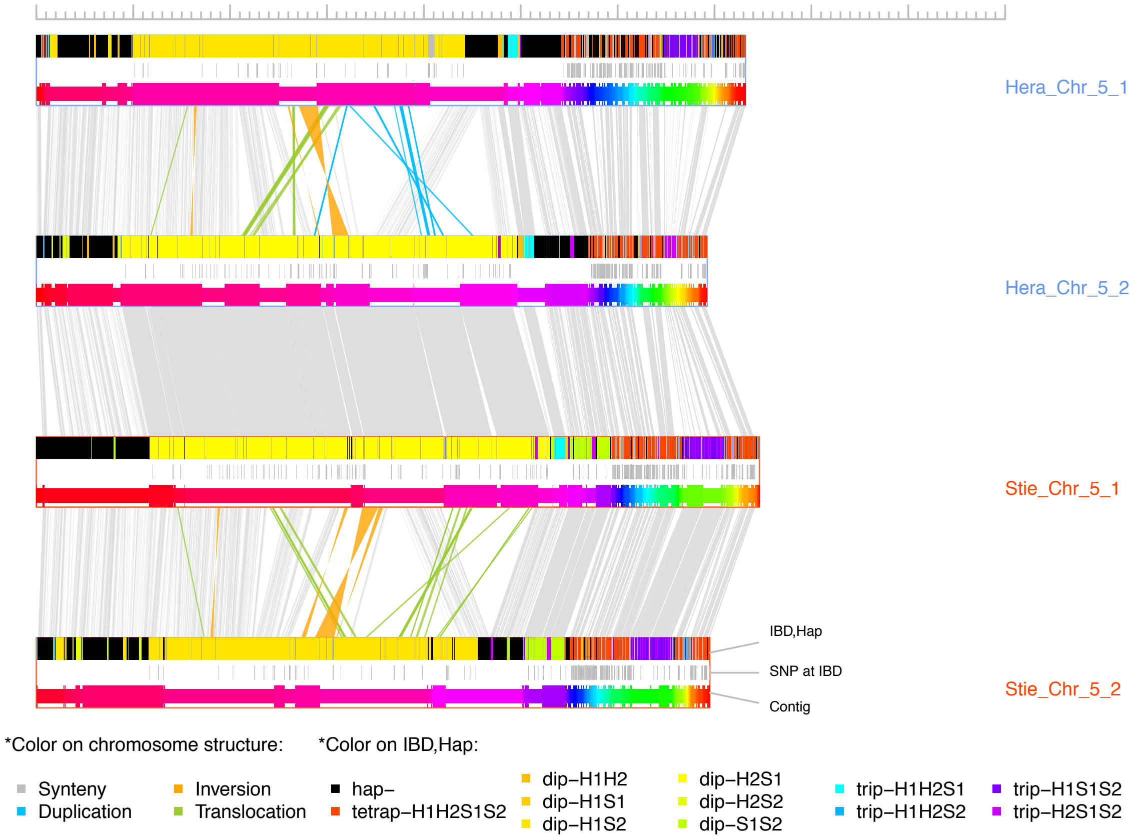


**Figure S15. Comparison of LG-wise haplotypes at LG 5**. Each of the four horizontal rows represents a haplotype of the chromosome (*Hera* 1, *Hera* 2, *Stieglitz* 1 and *Stieglitz* 2). Three types of information are within each haplotype box, top: IBD and unique regions along the chromosome (at 50 kb resolution), middle: distribution of SNPs at IBD regions, and bottom: scaffolded contigs. For each chromosome, three pair-wise structural comparisons are shown highlighting inversions (orange), duplications (blue), translocations (green) and syntenic regions (grey). Note that all the regions involving the breakpoints of the SVs ended within contigs. IBD regions can be shared by two (dip), three (trip) or even four (tetrap) haplotypes. The x-axis scale: 0-100 Mb.

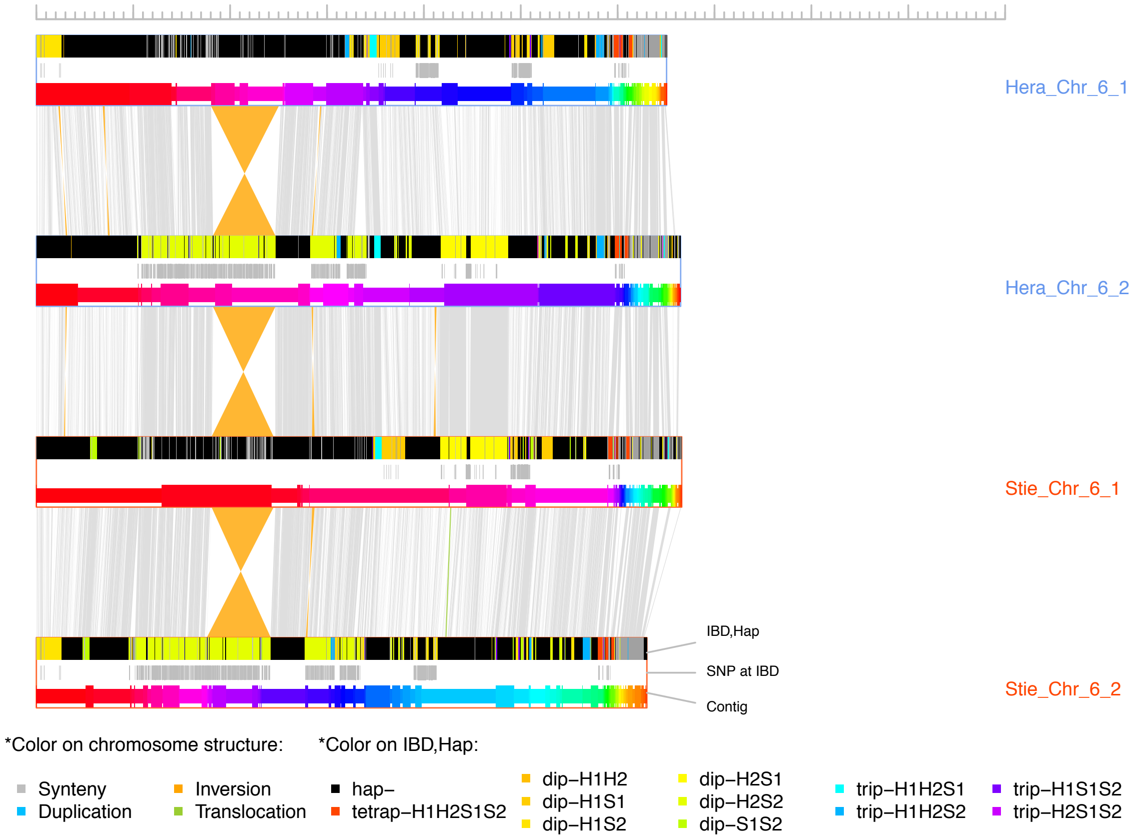


**Figure S16. Comparison of LG-wise haplotypes at LG 6**. Each of the four horizontal rows represents a haplotype of the chromosome (*Hera* 1, *Hera* 2, *Stieglitz* 1 and *Stieglitz* 2). Three types of information are within each haplotype box, top: IBD and unique regions along the chromosome (at 50 kb resolution), middle: distribution of SNPs at IBD regions, and bottom: scaffolded contigs. For each chromosome, three pair-wise structural comparisons are shown highlighting inversions (orange), duplications (blue), translocations (green) and syntenic regions (grey). Note that all the regions involving the breakpoints of the SVs ended within contigs. IBD regions can be shared by two (dip), three (trip) or even four (tetrap) haplotypes. The x-axis scale: 0-100 Mb.

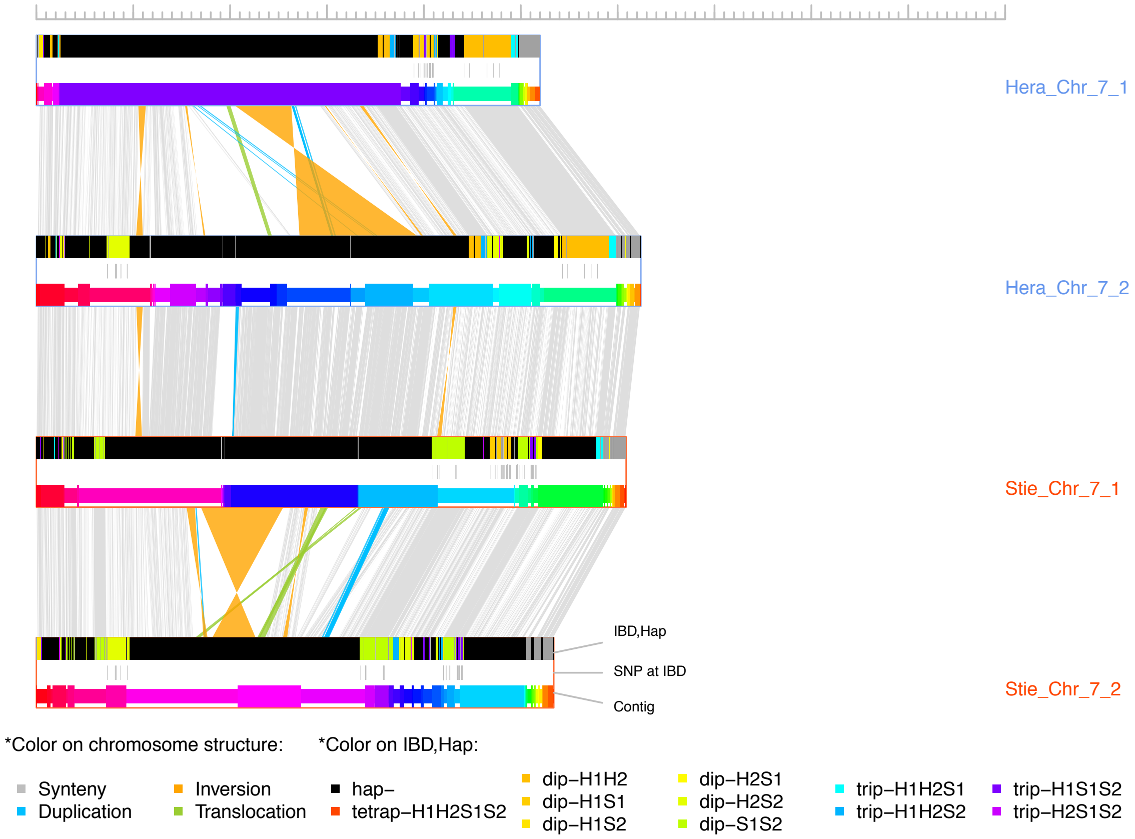


**Figure S17. Comparison of LG-wise haplotypes at LG 7**. Each of the four horizontal rows represents a haplotype of the chromosome (*Hera* 1, *Hera* 2, *Stieglitz* 1 and *Stieglitz* 2). Three types of information are within each haplotype box, top: IBD and unique regions along the chromosome (at 50 kb resolution), middle: distribution of SNPs at IBD regions, and bottom: scaffolded contigs. For each chromosome, three pair-wise structural comparisons are shown highlighting inversions (orange), duplications (blue), translocations (green) and syntenic regions (grey). Note that all the regions involving the breakpoints of the SVs ended within contigs. IBD regions can be shared by two (dip), three (trip) or even four (tetrap) haplotypes. The x-axis scale: 0-100 Mb.

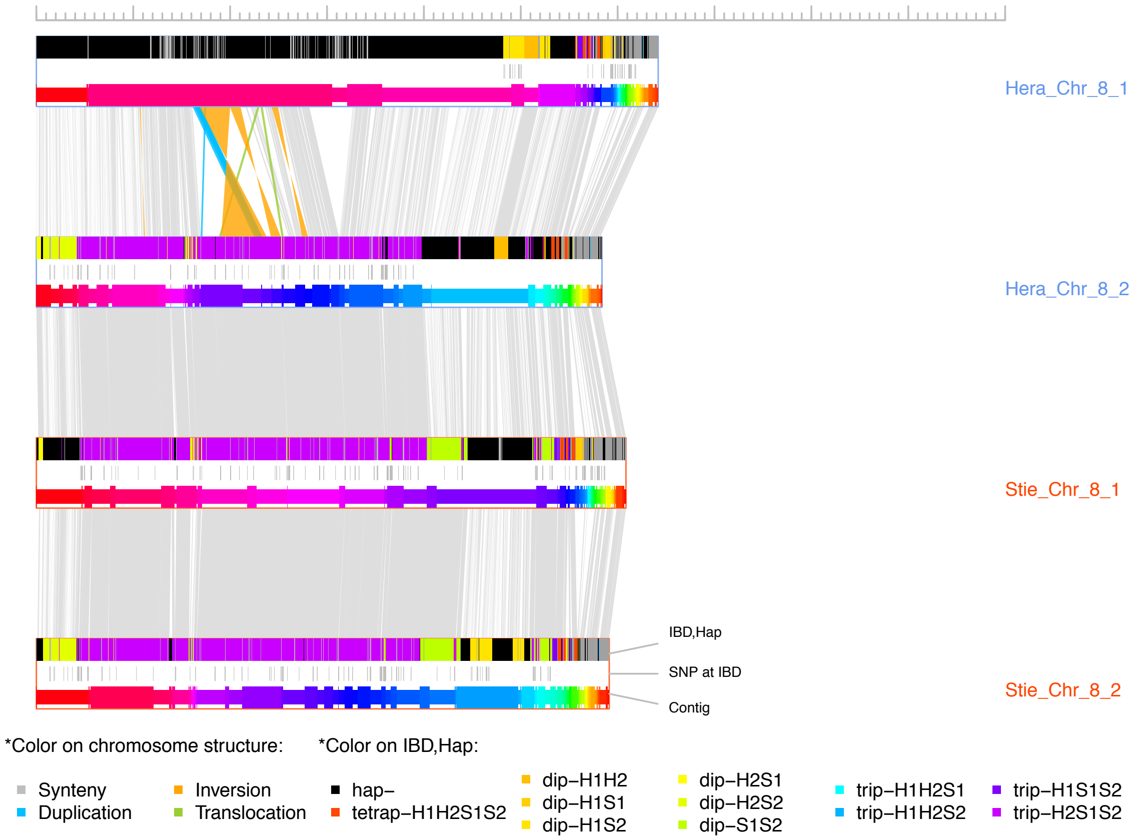


**Figure S18. Comparison of LG-wise haplotypes at LG 8**. Each of the four horizontal rows represents a haplotype of the chromosome (*Hera* 1, *Hera* 2, *Stieglitz* 1 and *Stieglitz* 2). Three types of information are within each haplotype box, top: IBD and unique regions along the chromosome (at 50 kb resolution), middle: distribution of SNPs at IBD regions, and bottom: scaffolded contigs. For each chromosome, three pair-wise structural comparisons are shown highlighting inversions (orange), duplications (blue), translocations (green) and syntenic regions (grey). Note that all the regions involving the breakpoints of the SVs ended within contigs. IBD regions can be shared by two (dip), three (trip) or even four (tetrap) haplotypes. The x-axis scale: 0-100 Mb.

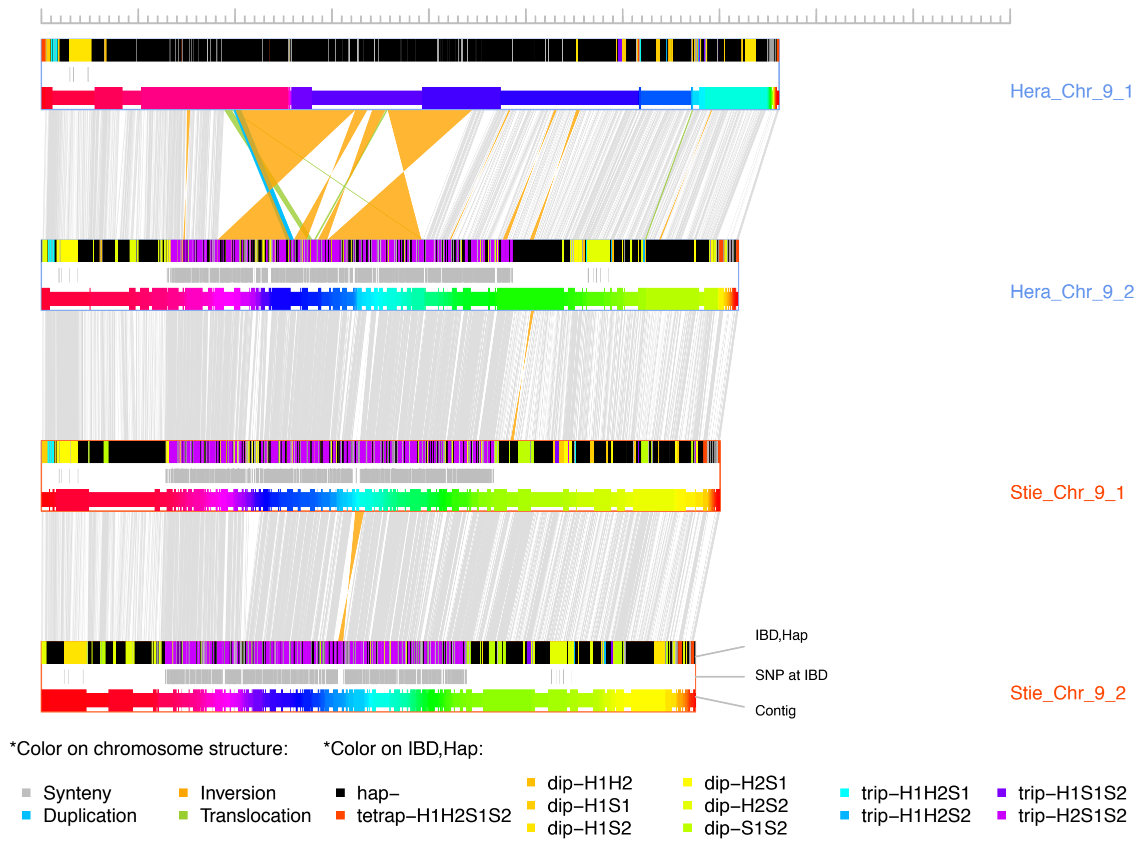


**Figure S19. Comparison of LG-wise haplotypes at LG 9**. Each of the four horizontal rows represents a haplotype of the chromosome (*Hera* 1, *Hera* 2, *Stieglitz* 1 and *Stieglitz* 2). Three types of information are within each haplotype box, top: IBD and unique regions along the chromosome (at 50 kb resolution), middle: distribution of SNPs at IBD regions, and bottom: scaffolded contigs. For each chromosome, three pair-wise structural comparisons are shown highlighting inversions (orange), duplications (blue), translocations (green) and syntenic regions (grey). Note that all the regions involving the breakpoints of the SVs ended within contigs. IBD regions can be shared by two (dip), three (trip) or even four (tetrap) haplotypes. The x-axis scale: 0-100 Mb.

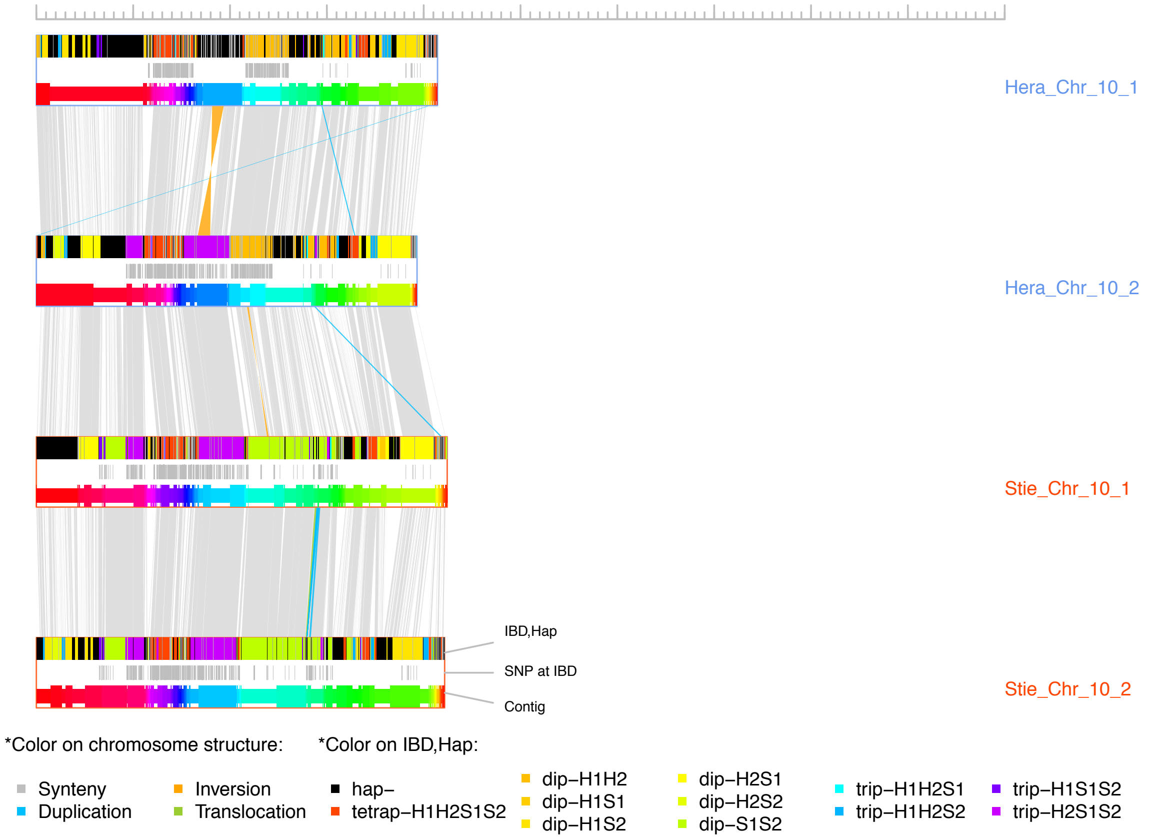


**Figure S20. Comparison of LG-wise haplotypes at LG 10**. Each of the four horizontal rows represents a haplotype of the chromosome (*Hera* 1, *Hera* 2, *Stieglitz* 1 and *Stieglitz* 2). Three types of information are within each haplotype box, top: IBD and unique regions along the chromosome (at 50 kb resolution), middle: distribution of SNPs at IBD regions, and bottom: scaffolded contigs. For each chromosome, three pair-wise structural comparisons are shown highlighting inversions (orange), duplications (blue), translocations (green) and syntenic regions (grey). Note that all the regions involving the breakpoints of the SVs ended within contigs. IBD regions can be shared by two (dip), three (trip) or even four (tetrap) haplotypes. The x-axis scale: 0-100 Mb.

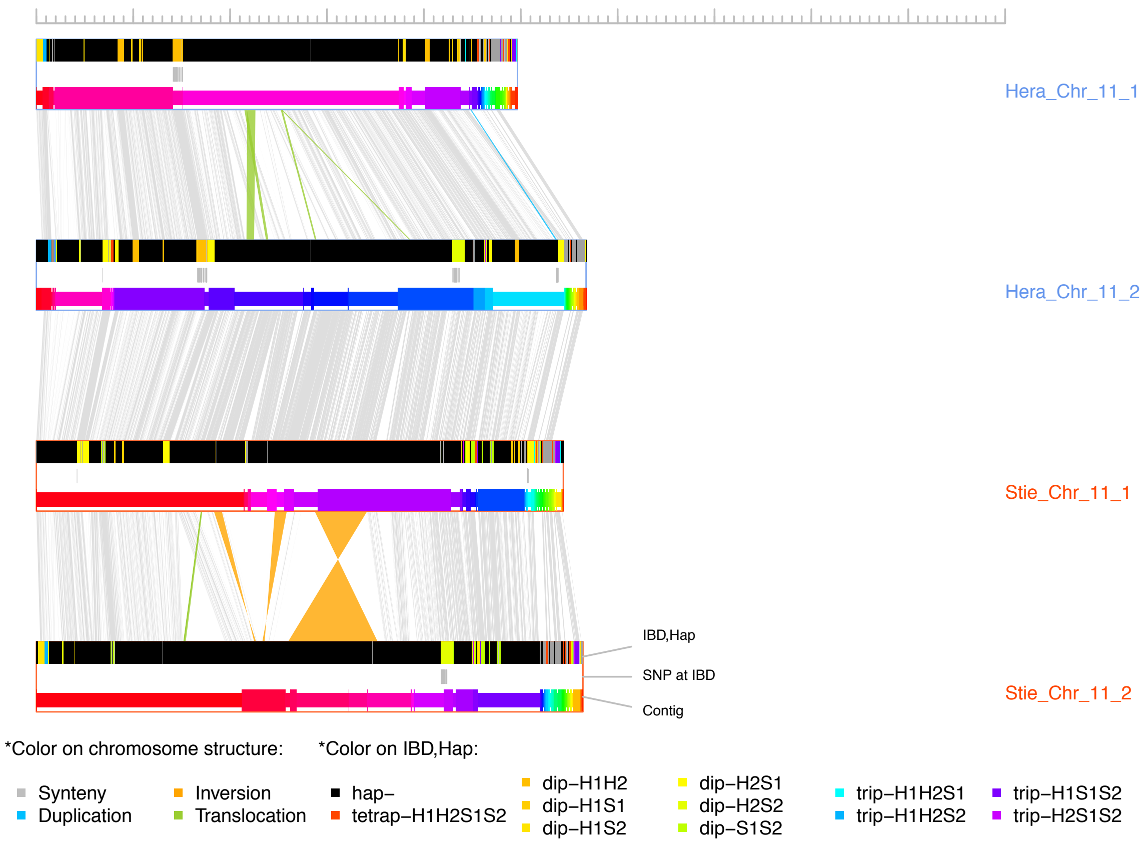


**Figure S21. Comparison of LG-wise haplotypes at LG 11**. Each of the four horizontal rows represents a haplotype of the chromosome (*Hera* 1, *Hera* 2, *Stieglitz* 1 and *Stieglitz* 2). Three types of information are within each haplotype box, top: IBD and unique regions along the chromosome (at 50 kb resolution), middle: distribution of SNPs at IBD regions, and bottom: scaffolded contigs. For each chromosome, three pair-wise structural comparisons are shown highlighting inversions (orange), duplications (blue), translocations (green) and syntenic regions (grey). Note that all the regions involving the breakpoints of the SVs ended within contigs. IBD regions can be shared by two (dip), three (trip) or even four (tetrap) haplotypes. The x-axis scale: 0-100 Mb.

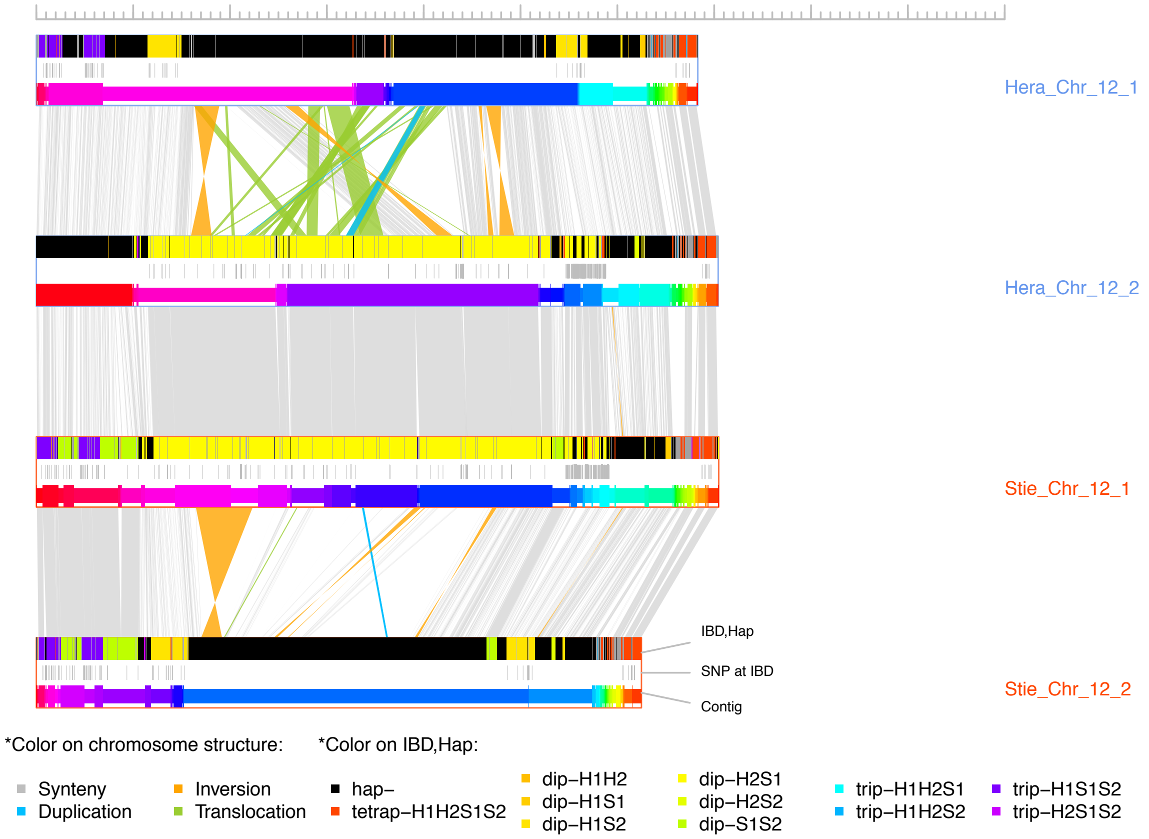


**Figure S22. Comparison of LG-wise haplotypes at LG 12**. Each of the four horizontal rows represents a haplotype of the chromosome (*Hera* 1, *Hera* 2, *Stieglitz* 1 and *Stieglitz* 2). Three types of information are within each haplotype box, top: IBD and unique regions along the chromosome (at 50 kb resolution), middle: distribution of SNPs at IBD regions, and bottom: scaffolded contigs. For each chromosome, three pair-wise structural comparisons are shown highlighting inversions (orange), duplications (blue), translocations (green) and syntenic regions (grey). Note that all the regions involving the breakpoints of the SVs ended within contigs. IBD regions can be shared by two (dip), three (trip) or even four (tetrap) haplotypes. The x-axis scale: 0-100 Mb.

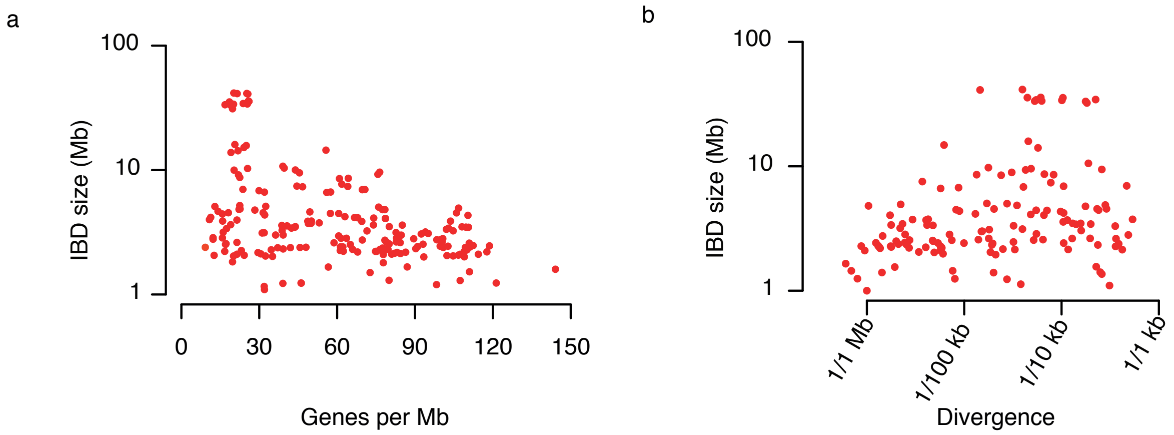


**Figure S23. Correlation of features at IBD blocks**. a. Correlation of IBD block size with gene density. IBD blocks could reach over 40 Mb, however, such large IBD blocks located in peri-centromeric regions were usually with low gene density. b. Correlation of IBD block size with their age estimated based on the accumulated mutations. IBD blocks showed different levels of mutations indicating that larger IBD blocks were generally not younger than many of the smaller IBD blocks. Y-axis was in log10 scaled.

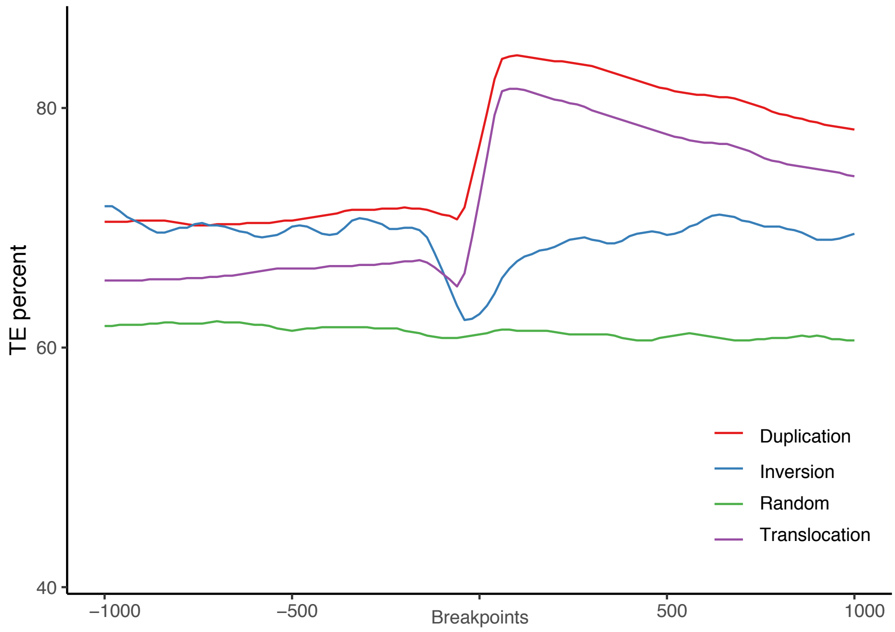


**Figure S24. Enrichment of transposon elements in structural variations**. TE percent is much higher in duplications and translocations, but not in inversions.

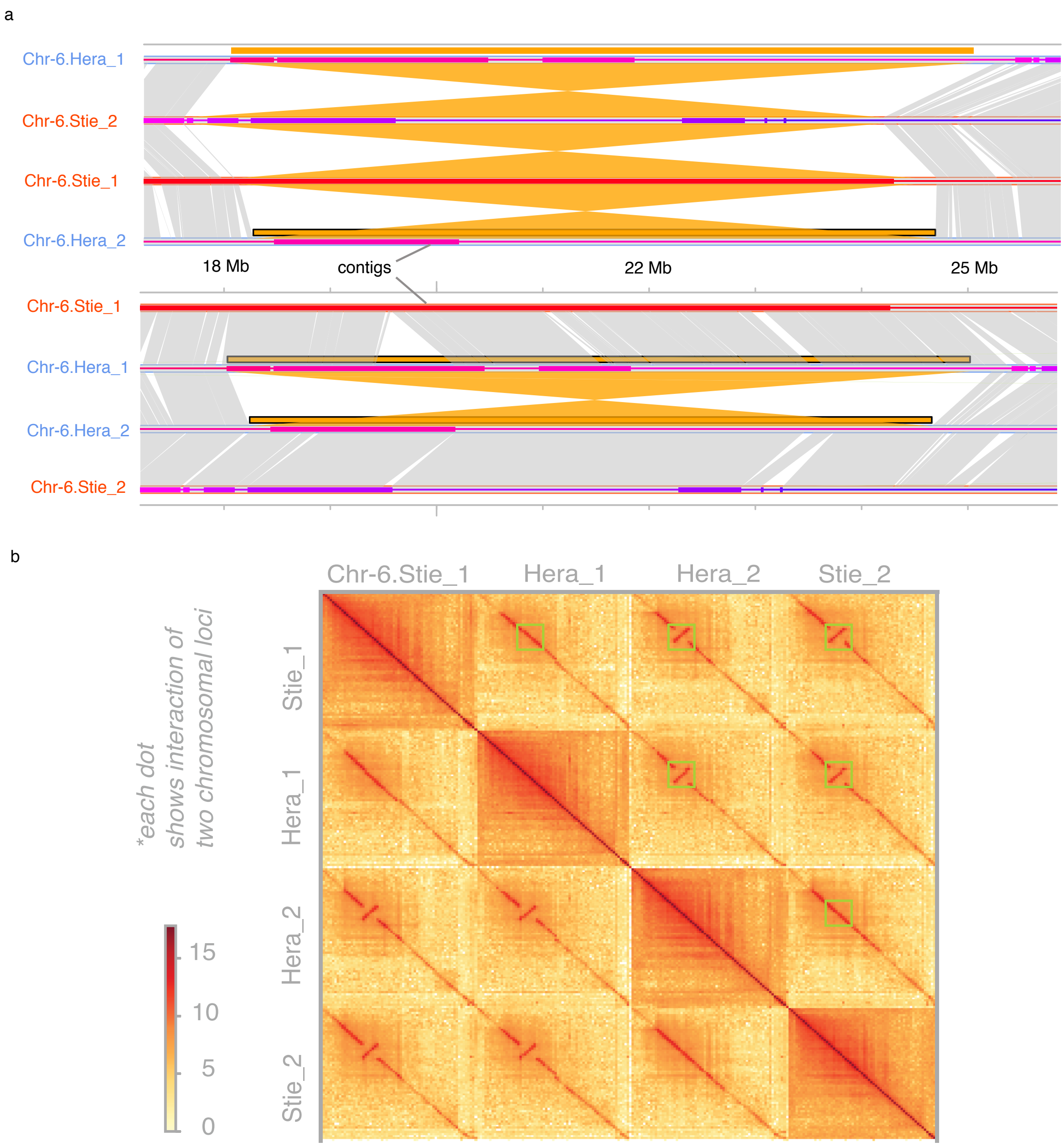


**Figure S25. Example for validation of structural variation. a**: Pairwise comparison of the four haplotypes of Chr. 6 revealed four inversions:

*Chr-6.Hera_1:18,032,256-25,020,440* versus *Chr-6.Stie_2:17,620,164-24,170,977*,
*Chr-6.Stie_2:17,620,163-24,170,977* versus *Chr-6.Stie_1:18,104,687-24,467,329*,
*Chr-6.Stie_1:18,104,687-24,467,329* versus *Chr-6.Hera_2:18,241,689-24,660,203*,
*Chr-6.Hera_1:18,032,256-25,020,440* versus *Chr-6.Hera_2:18,241,690-24,660,203*.

Two other pairwise comparisons, i.e., *Chr-6.Stie_1* and *Chr-6.Hera_1, Chr-6.Hera_2* and *Chr-6.Stie_2*, showed high levels of synteny in the respective regions. All the regions involving the breakpoints of the inversions ended within contigs. **b**. The same inversions/synteny relationship between haplotypes were observed in Hi-C contact map. Together with information from **a**, these data evidenced that the inversions are real.

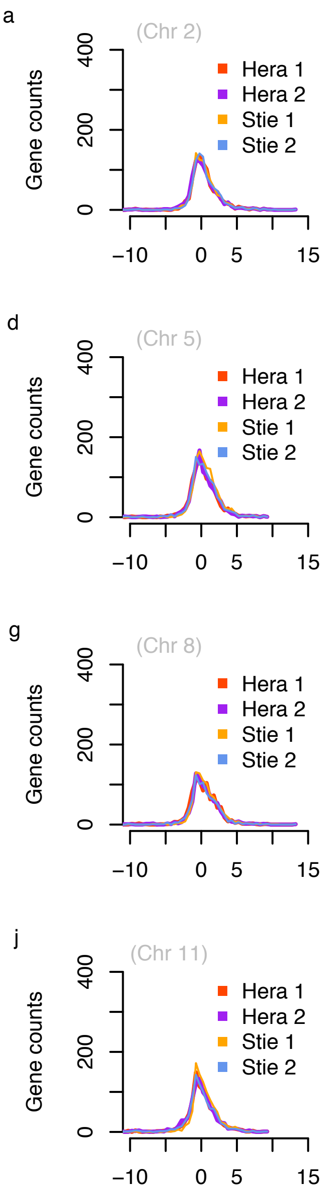


**Figure S26. Four haplotypes of chromosome 2-12 showed comparable amount of gene expression** (FPKM: fragments per kilobase per million reads). Note, chromosome 1 was given in main text Fig. 4d.

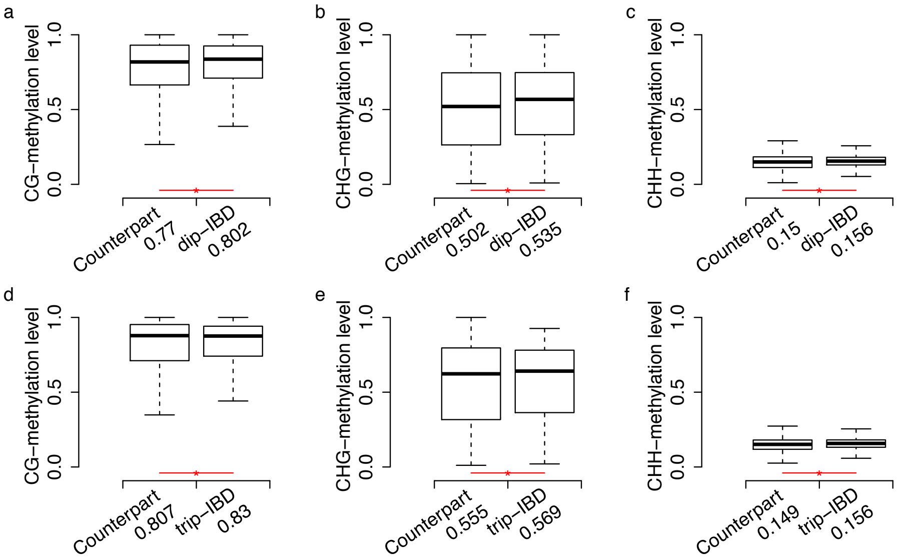


**Figure S27. Comparison of methylation (at CG, CHG, CHH context) between IBD blocks (of 50 kb) and their counterparts (i.e., genomic regions in synteny with related IBDs).** a-c. Level of CG/CHG/CHH methylation related to IBD blocks shared by two haplotypes (labeled with "dip-IBD"). d-f. Level of CG/CHG/CHH methylation related to IBD blocks shared by three haplotypes (labeled with "trip-IBD"). Mean values of methylation levels (Materials and Methods) among the investigated blocks are given after *x*-axis labels. The number of analyzed dip-IBD blocks was 4,706, and that of dip-IBD counterpart blocks was also 4,706. Number of analyzed trip-IBD blocks was 2,925, and that of trip-IBD counterpart blocks was 975. The red line with an asterisk under each pair of boxes indicates that there is a significant difference between the two sets (*t*-test, *p*-value<0.05). In general, the methylation level at IBD blocks were slightly but significantly higher than that at their non-IBD counterparts. For example, in **a**, the average CG-methylation level of the 4,706 dip-IBD blocks was 0.802 and the methylation level of their counterparts was 0.77, while the former was siginicantly higher than the latter.

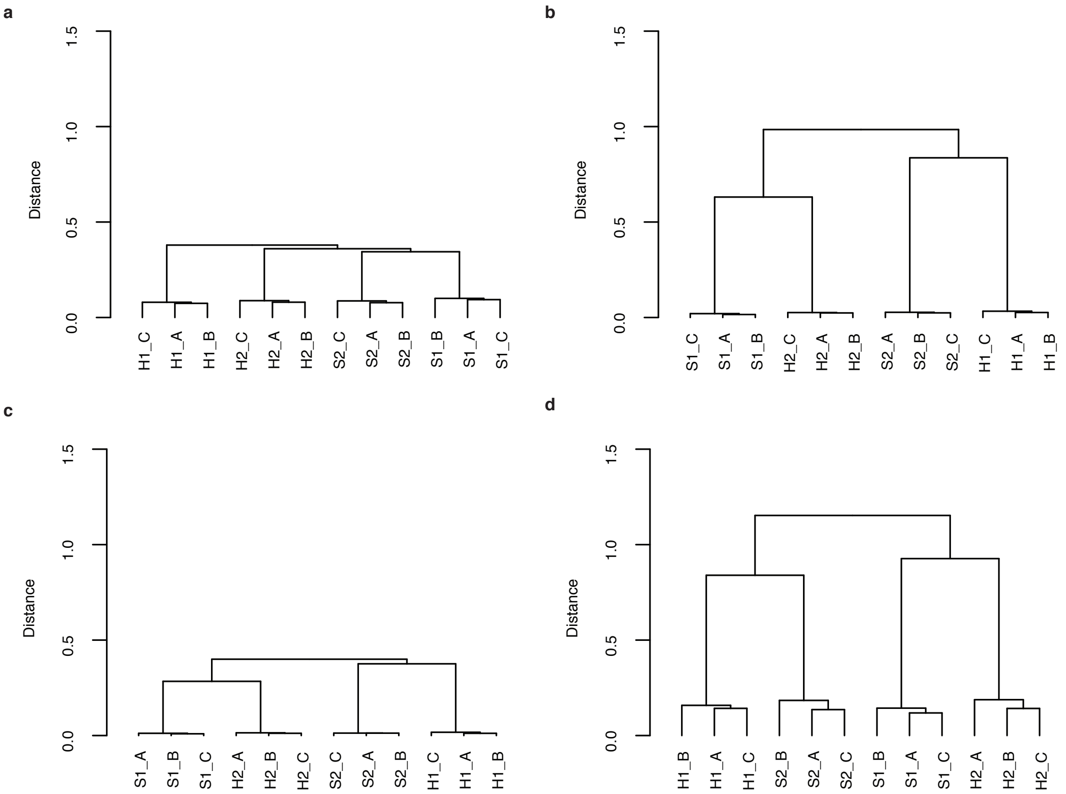


**Figure S28. High consistency between the technical replicates of sequencings of gene expression or DNA methylation as shown by well-clustered haplotypes. a.** Clustering of replicates regarding haplotypes using haplotype-specific genome-wide allele expression. **b.** Clustering of replicates regarding haplotypes using haplotype-specific genome-wide methylation in CG context. **c.** Clustering of replicates regarding haplotypes using haplotype-specific genome-wide methylation in CHG context. **d.** Clustering of replicates regarding haplotypes using haplotype-specific genome-wide methylation in CHH context.

#### Supplementary Reference

1. J.A. Campoy, H. Sun *et al*. *Gamete binning*: chromosome-level and haplotype-resolved genome assembly enabled by high-throughput single-cell sequencing of gamete genomes. *Genome Biol.* **21**, 306 (2020).
2. M. Marçais, C. Kingsford. A fast, lock-free approach for efficient parallel counting of occurrences of k-mers. *Bioinformatics* **27**, 764-770 (2011).
3. H. Sun, J. Ding, M. Piednoël, K. Schneeberger. *FindGSE*: Estimating genome size variation within human and *Arabidopsis* using *k*-mer frequencies. *Bioinformatics* **34**, 550-557 (2018).
4. H. Cheng, G.T. Concepcion, X. Feng, H. Zhang, H. Li. Haplotype-resolved de novo assembly with phased assembly graphs. *Nat Methods* **18**, 170-175 (2021).
5. B. Langmead, S.L. Salzberg. Fast gapped-read alignment with *Bowtie* 2. *Nature methods* **9**, 357-359 (2012).
6. B.J. Walker et al. *Pilon*: An integrated tool for comprehensive microbial variant detection and genome assembly improvement. *PLoS One* **9**, 1-14 (2014).
7. H. Li. *Minimap2*: Pairwise alignment for nucleotide sequences. *Bioinformatics* **34**(18), 3094-100 (2018).
8. H. Li. et al. The Sequence Alignment/Map format and SAMtools. *Bioinformatics* **25**, 2078-2079 (2009).
9. The Potato Genome Sequencing Consortium. Genome sequence and analysis of the tuber crop potato. *Nature* **475**, 189-195 (2011).
10. G.M. Pham, J.P. Hamilton *et al*. Construction of a chromosome-scale long-read reference genome assembly for potato. *GigaScience* **9**, 1-11 (2020).
11. A.R. Quinlan, I.M. Hall. *BEDTools*: A flexible suite of utilities for comparing genomic features. *Bioinformatics* **26**, 841-842 (2010).
12. R. Vaser, I. Sović, N. Nagarajan, M. Šikić. (2017). Fast and accurate de novo genome assembly from long uncorrected reads. *Genome Res*. **27**, 737-746.
13. S.F. Altschul, W. Gish, W. Miller, E.W. Myers & D.J. Lipman. Basic local alignment search

tool. *J. Mol. Biol.* **215**, 403-410 (1990).

14. M. Kokot, M. Dlugosz, S. Deorowicz. *KMC3*: counting and manipulating *k*-mer statistics. *Bioinformatics* **33**, 2759-2761 (2017).
15. H. Li, R. Durbin. Fast and accurate short read alignment with Burrows-Wheeler transform. *Bioinformatics* **25**, 1754-1760 (2009).
16. X. Zhang, S. Zhang, Q. Zhao, R. Ming, H. Tang. Assembly of allele-aware, chromosomal-scale autopolyploid genomes based on Hi-C data. *Nat. Plants* **5**, 833-845 (2019).
17. J. Ghurye, et al. Integrating Hi-C links with assembly graphs for chromosome-scale assembly. *PLoS Comput. Biol*. **15**, 1-19 (2019).
18. A. Rhie, B.P. Walenz, S. Koren, A.M. Phillippy. *Merqury*: reference-free quality, completeness, and phasing assessment for genome assemblies. *Genome Biol* **21**, 1-27 (2020).
19. M. Goel, H. Sun, W.B. Jiao, K. Schneeberger. *SyRI*: finding genomic rearrangements and local sequence differences from whole-genome assemblies. *Genome Biol.* **20**, 1-13 (2019).
20. M. Stanke, *et al*. *AUGUSTUS*: *Ab initio* prediction of alternative transcripts. *Nucleic Acids Res.* **34**, 435-439 (2006).
21. W.H. Majoros, M. Pertea, S.L. Salzberg. *TigrScan and GlimmerHMM*: Two open source ab initio eukaryotic gene-finders. *Bioinformatics* **20**, 2878-2879 (2004).
22. A.D. Johnson *et al*. *SNAP*: A web-based tool for identification and annotation of proxy SNPs using HapMap. *Bioinformatics* **24**, 2938-2939 (2008).
23. Q. Zhou, D. Tang *et al*. Haplotype-resolved genome analyses of a heterozygous diploid potato. *Nat. Genet.* **52**, 1018-1023 (2020).
24. G.S.C. Slater, E. Birney. Automated generation of heuristics for biological sequence comparison. *BMC Bioinformatics* **6**, 1-11 (2005).
25. D. Kim. *et al*. Graph-based genome alignment and genotyping with *HISAT*2 and *HISAT*-genotype. *Nat. Biotechnol*. **37**, 907-915 (2019).
26. M. Pertea. et al. *StringTie* enables improved reconstruction of a transcriptome from RNA-seq reads. *Nat. Biotechnol.* **33**, 290-295 (2015).

27. B. Haas, B. *et al.* Automated eukaryotic gene structure annotation using *EVidenceModeler* and the Program to Assemble Spliced Alignments. *Genome Biol.* **9**, 1-22 (2008).
28. O. Keller, F. Odronitz, M. Stanke, M. Kollmar, S. Waack. *Scipio*: Using protein sequences to determine the precise exon/intron structures of genes and their orthologs in closely related species. *BMC Bioinformatics* **9**, 1-12 (2008).
29. F.A. Simão, R.M. Waterhouse *et al*. *BUSCO*: Assessing genome assembly and annotation completeness with single-copy orthologs. *Bioinformatics* **31**, 3210-3212 (2015).
30. P. Jones *et al*. *InterProScan* 5: genome-scale protein function classification. *Bioinformatics* **30**, 1236-1240 (2014).
31. G. Yu, L. Wang, Y. Han, Q. He. *clusterProfiler*: an R package for comparing biological themes among gene clusters. *OMICS: A Journal of Integrative Biology* **16**, 284-287 (2012).
32. E.P. Nawrocki, S.R. Eddy. *Infernal 1.1*: 100-fold faster RNA homology searches. *Bioinformatics* **29**, 2933-2935 (2013).
33. I. Kalvari, E.P. Nawrocki, N. Ontiveros-Palacios *et al*. *Rfam 14*: expanded coverage of metagenomic, viral and microRNA families. *Nucleic Acids Res*. 49, D192-D200 (2021).
34. V. Wucher, F. Legeai, B. Hédan, G. Rizk, L. Lagoutte L et al. *FEELnc*: a tool for long non-coding RNA annotation and its application to the dog transcriptome. *Nucleic Acids Res.*, **45**, e57 (2017).
35. Y.J. Kang, D.C. Yang, L. Kong, M. Hou, Y.Q. Meng, L. Wei, G. Gao. CPC2: a fast and accurate coding potential calculator based on sequence intrinsic features. *Nucleic Acids Res.* **45**, W12-W16 (2017).
36. UniProt Consortium. Activities at the Universal Protein Resource (UniProt). *Nucleic Acids Res.* **42**, D191-8 (2014).
37. X. Zhao, J. Li, B. Lian, H. Gu, Y. Li, Y. Qi. Global identification of Arabidopsis lncRNAs reveals the regulation of MAF4 by a natural antisense RNA. *Nat Commun* **9**, 5056 (2018).
38. D.M. Emms, S. Kelly. *OrthoFinder*: Phylogenetic orthology inference for comparative genomics. *Genome Biol.* **20**, 1-14 (2019).
39. X. Hu *et al*. *pIRS*: Profile-based Illumina pair-end reads simulator. *Bioinformatics* **28**, 1533-

1535 (2012).

40. A.M. Bolger, M. Lohse, B. Usadel. *Trimmomatic*: a flexible trimmer for Illumina sequence data. *Bioinformatics* **30**, 2114-2120 (2014).

41. S. Anders, P.T. Pyl, W. Huber. *HTSeq*--a Python framework to work with high-throughput sequencing data. *Bioinformatics* **15**, 166-9 (2015).

42. C.W. Law, M. Alhamdoosh, S. Su, X. Dong, L. Tian et al. RNA-seq analysis is easy as 1-2-3 with limma, Glimma and edgeR. *F1000Research* **5**, ISCB Comm J-1408 (2016).

43. F. Krueger, S.R. Andrews. *Bismark*: a flexible aligner and methylation caller for Bisulfite-Seq applications. *Bioinformatics* **27**, 1571-2 (2011).

44. R.C.B. Hutten and R.van Berloo. An online potato pedigree database. URL: http://www.plantbreeding.wur.nl/PotatoPedigree/ (2001).

45. R.van Berloo, R.C.B. Hutten, H.J.van Eck, and R.G.F Visser. An online potato pedigree database resource. *Potato research* **50**, 45-57 (2007).

**Data S1** (Data_S1.xlsx)**:** Overview of sequencing data

**Data S2** (Data_S2.xlsx) **:** Gene annotation in Otava (and comparison to DM and RH)

**Data S3** (Data_S3.xlsx)**:** Summary on Benchmarking Universal Single Copy Orthologs (BUSCO) evaluation for assemblies and annotations (with comparisons to existing assemblies)

**Data S4** (Data_S4.xlsx): Noncoding RNA

**Data S5** (Data_S5.xlsx)**:** Repeat statistics

**Data S6** (Data_S6.xlsx)**:** Statistics on rDNA

**Data S7** (Data_S7.xlsx)**:** Potentially collapsed 18,534 SNPs at IBD regions

**Data S8** (Data_S8.xlsx): Enrichment of transposon elements (TE) among structural variations (SV)

**Data S9** (Data_S9.xlsx)**:** Large structural variations between the four haplotypes of each chromosome

**Data S10** (Data_S10.xlsx)**:** PAVs in *R* Genes

**Data S11** (Data_S11.xlsx): Statistics of 1219 differentially expressed genes

**Data S12** (Data_S12.xlsx): Statistics on correlation of methylation and expression of 304 genes
